# Mitochondrial leakage and mtDNA damage trigger early immune response in Inclusion Body Myositis

**DOI:** 10.1101/2024.08.05.606624

**Authors:** Felix Kleefeld, Emily Cross, Daniel Lagos, Benedikt Schoser, Andreas Hentschel, Tobias Ruck, Christopher Nelke, Sara Walli, Katrin Hahn, Denisa Hathazi, Andrew L. Mammen, Maria Casal-Dominguez, Marta Gut, Ivo Glynne Gut, Simon Heath, Anne Schänzer, Hans-Hilmar Goebel, Iago Pinal-Fernandez, Andreas Roos, Corinna Preuße, Werner Stenzel, Rita Horvath

## Abstract

Polymyositis with mitochondrial pathology (PM-Mito) was first identified in 1997 as a subtype of idiopathic inflammatory myopathy. Recent findings demonstrated significant molecular similarities between PM-Mito and Inclusion Body Myositis (IBM), suggesting a trajectory from early to late IBM and prompting the inclusion of PM-Mito as an IBM precursor (early IBM) within the IBM spectrum. Both PM-Mito and IBM show mitochondrial abnormalities, suggesting mitochondrial disturbance is a critical element of IBM pathogenesis.

The primary objective of this cross-sectional study was to characterize the mitochondrial phenotype in PM-Mito at histological, ultrastructural, and molecular levels and to study the interplay between mitochondrial dysfunction and inflammation. Skeletal muscle biopsies of 27 patients with PM-Mito and 27 with typical IBM were included for morphological and ultrastructural analysis. Mitochondrial DNA (mtDNA) copy number and deletions were assessed by qPCR and long-range PCR, respectively. In addition, full-length single-molecule sequencing of the mtDNA enabled precise mapping of deletions. Protein and RNA levels were studied using unbiased proteomic profiling, immunoblotting, and bulk RNA sequencing. Cell-free mtDNA (cfmtDNA) was measured in the serum of IBM patients.

We found widespread mitochondrial abnormalities in both PM-Mito and IBM, illustrated by elevated numbers of COX-negative and SDH-positive fibers and prominent ultrastructural abnormalities with disorganized and concentric cristae within enlarged and dysmorphic mitochondria. MtDNA copy numbers were significantly reduced, and multiple large-scale mtDNA deletions were already evident in PM-Mito, compared to healthy age-matched controls, similar to the IBM group. The activation of the canonical cGAS/STING inflammatory pathway, possibly triggered by the intracellular leakage of mitochondrial DNA, was evident in PM-Mito and IBM. Elevated levels of circulating cfmtDNA also indicated leakage of mtDNA as a likely inflammatory trigger. In PM-Mito and IBM, these findings were accompanied by dysregulation of proteins and transcripts linked to the mitochondrial membranes.

In summary, we identified that mitochondrial dysfunction with multiple mtDNA deletions and depletion, disturbed mitochondrial ultrastructure, and defects of the inner mitochondrial membrane are features of PM-Mito and IBM, underlining the concept of an IBM-spectrum disease (IBM-SD). The activation of inflammatory pathways related to mtDNA release indicates a significant role of mitochondria-associated inflammation in the pathogenesis of IBM-SD. Thus, mitochondrial abnormalities precede tissue remodeling and infiltration by specific T-cell subpopulations (e.g., KLRG1^+^) characteristic of late IBM. This study highlights the critical role of early mitochondrial abnormalities in the pathomechanism of IBM, which may lead to new approaches to therapy.

## Introduction

Polymyositis with mitochondrial pathology (PM-Mito), first identified in 1997 as a subtype of idiopathic inflammatory myopathy (IIM), is characterized by significant mitochondrial abnormalities and inflammation.^1^ Recently, we described a considerable histological and molecular overlap between PM-Mito and Inclusion Body Myositis (IBM).^2^ We showed that >90% of patients diagnosed with PM-Mito developed IBM in their later course, which was confirmed by follow-up biopsies several years apart, and proposed to include PM-Mito as an early form of IBM (eIBM) within the spectrum of IBM (IBM spectrum disease, IBM-SD).^2^ While specific inflammatory features and characteristic tissue damage of skeletal muscle, including protein aggregates and rimmed vacuoles, are less pronounced in PM-Mito compared to typical IBM, we observed a similar quantity and quality of mitochondrial abnormalities defined by light microscopy (COX-SDH-positive fibers), pointing towards mitochondrial abnormalities and likely dysfunction being an early feature of IBM. Indeed, multiple mtDNA deletions, typically seen in IBM, have also been reported in smaller cohorts of PM-Mito patients.^3^ However, mitochondrial abnormalities in PM-Mito have not been studied in depth. Mitochondrial morphological abnormalities with multiple mtDNA deletions and depletion were shown as a characteristic feature of IBM.^4,5^ However, the mechanisms linking these mitochondrial abnormalities and inflammation to the pathogenesis of IBM remained puzzling.

Recently, abnormal mitochondria have been reported to contribute to interferon-mediated inflammation in dermatomyositis by activating the canonical cGAS/STING (cyclic GMP-AMP synthase/stimulator of interferon genes) pathway.^6^ CGAS/STING activation has also been identified in other IIM, including immune-mediated necrotizing myopathy (IMNM)^7^ and in a range of autoinflammatory syndromes such as hereditary interferonopathies.^8^ Notably, the role of the cGAS/STING pathway in IBM remains unknown to date.

In IBM, a link has been suggested between degenerative features (e.g., TDP-43 deposition) and mitochondrial pathology.^9^ These findings have led to the assumption that IBM might be a primary degenerative disease of skeletal muscle.^10^ Moreover, this hypothesis has been supported by the failure of previous immunosuppressive treatment strategies. However, IBM also shows distinct features of a complex immune-mediated disorder, possibly involving cN1A (cytosolic 5’-nucleotidase 1A) antibodies, CD8+ lymphocytes, macrophages, and cytokines such as interferons (IFN).^11–13^ Yet, immunization with cN1A antibodies could only reproduce inflammatory features but no mitochondrial abnormalities in mice.^14^ Despite extensive research, whether mitochondrial dysfunction should be viewed as a downstream consequence of persistent inflammation or mitochondria might be at the core of disease pathophysiology remains unsolved.

In the present study, we hypothesized that mitochondrial dysfunction is an early feature of IBM and might be mechanistically linked to the development of autoimmunity and inflammation. Studying a well-characterized, unique cohort of PM-Mito muscle biopsy specimens, our primary research focused on providing an in-depth characterization of the mitochondrial phenotype observed in PM-Mito at the histological, ultrastructural, and molecular levels. In addition, we investigated the interplay between early mitochondrial damage and activation of canonical inflammatory pathways. We identified that mitochondrial dysfunction, including multiple mtDNA deletions and mtDNA depletion, along with disturbed mitochondrial ultrastructure and membrane defects, is an early feature of IBM. Activation of inflammatory pathways related to mtDNA release highlights the significant role of mitochondria-associated inflammation in IBM pathogenesis, where mitochondrial abnormalities precede tissue remodeling and T-cell infiltration.

## Materials and methods

For this study, experiments were conducted in Berlin, Germany (Charité – Universitätsmedizin Berlin, Dep. of Neuropathology; histology and electron microscopy), Cambridge, United Kingdom (University of Cambridge, Dep. of Clinical Neurosciences; immunoblotting, long-range, and qPCR), Barcelona, Spain (Centro Nacional de Análisis Genómico (CNAG); nanopore sequencing of mtDNA), Dortmund, Germany (Leibniz-Institut für Analytische Wissenschaften – ISAS e.V.; unbiased proteomic profiling), and Bethesda, USA (the National Institutes of Health; bulk RNAseq). Approval for the study was obtained from the institutional ethics review board of Charité – Universitätsmedizin Berlin (EA2/107/14; EA1/019/22), and a Material Transfer Agreement (MTA) was established between the participating institutions. The study adhered to the principles outlined in the 1964 Declaration of Helsinki, and patient and control data were anonymized following the guidelines of the local institutional ethics review board.

### Patients, Samples, and Clinical Assessment

This study included skeletal muscle biopsy specimens showing signs of Polymyositis with mitochondrial pathology (referred to as ‘PM-Mito’) and clinically and pathologically confirmed IBM according to ENMC criteria.^15^ The definition of PM-Mito was based on established histopathological criteria, including endomysial inflammation, invasion of non-necrotic muscle fibers, and >3% of muscle fibers with deficient COX activity (visualized by combined COX-SDH enzyme histochemical preparations).^1^ Clinical and serological (auto-antibodies) features were not considered when classifying PM-Mito patients. All available biopsies were included with signs of PM-Mito from the cohorts in Berlin and Giessen, Germany. To minimize selection bias, IBM biopsies were randomly selected from available samples in Berlin without preselection based on comorbidities, gender, or antibody status. Control biopsy specimens from individuals with normal CK levels, negative myositis line blots (Myositis profile 4 *EUROLINE immunoblot;* EUROIMMUN AG, Lübeck, Germany), and unremarkable skeletal muscle morphology were also included. None of the controls had a family history of neuromuscular disease. All biopsies were obtained for diagnostic purposes at Charité-Universitätsmedizin Berlin, Germany, and Justus Liebig University Giessen, Germany, and analyzed by myopathologists (HHG, WS, FK, AS). All patients’ medical data, including a complete neurological examination at the time of biopsy, were collected. In addition, laboratory data (creatine kinase, CK; myositis line blot) were collected.

### Skeletal muscle biopsies

We examined skeletal muscle biopsy specimens from PM-Mito (n=27) and IBM (n=27) patients, all sourced from quadriceps femoris muscles. The specimens were snap-frozen and cryopreserved at -80°C before analysis.

### Histological and Immunohistochemical Analysis

Samples were processed following standardized protocols at the Department of Neuropathology, Charité - Universitätsmedizin, Berlin. Routine staining, immunohistochemical, and double immunofluorescence reactions were carried out as described previously.^2,16,17^ In short, the 7-µm-thick cryostat sections were stained with hematoxylin & eosin, modified Gömöri trichrome, for nonspecific esterase, alkaline phosphatase, etc., as described in a consensus statement^18^ and with a comprehensive panel of antibodies for immunohistochemistry and double immunofluorescence as described previously.^16,17^ Appropriate positive and negative controls (tissue reactions) were used where necessary. Additionally, histologically normal muscles (e.g., for MHC class I or II positivity of capillaries) were used as negative controls for immunohistochemical reactions. The above-mentioned comprehensive antibody panel was also used to ensure negative staining results by studying so-called “irrelevant antibodies” for validation. For quantification of mitochondrial morphological alterations (i.e., COX-negativity) and immune cells (CD8+ T-cells, CD68+ macrophages, CD45+ leukocytes), muscle fibers and immune cells were counted in ten high power fields (HPF, based on the microscope used and the respective oculars (Olympus WH10x-H/22) ≙ 0.16 mm^2^).

### Transmission electron microscopy

For transmission electron microscopy (TEM), muscle biopsy specimens were fixed and embedded according to standard protocols and as described previously.^19^ In brief, muscle specimens were fixed in 2.5% GA diluted in 0.1 M sodium cacodylate buffer for a minimum of 24 hours at four °C, osmicated in 1% osmium tetroxide in 0.05 M sodium cacodylate buffer, dehydrated using graded acetone series including combined *en-bloc* staining with 1% uranyl acetate and 0.1% phosphotungstic acid in 70% acetone, infiltrated in RenLam resin and then polymerized for 48–72 h at 60°C. Semithin sections (500 nm) were stained with Richardson solution (methylene blue) for microanatomical examination. Ultrathin sections (60–70 nm) were stained with uranyl acetate and lead citrate. Next, ultrastructural analysis was performed using TEM 902 and TEM 906 (Zeiss, Oberkochen, Germany).

### Unbiased proteomic studies

Muscle biopsy specimens were prepared for proteomic analysis following established protocols and as described previously.^20^ The analysis included 14 samples from PM-Mito patients and ten from IBM patients. Five adult male and two female patients with normal family histories, histological studies, and laboratory work-ups served as controls. These individuals were biopsied diagnostically for muscle pain but showed no pathological changes upon microscopic examination. Using a bottom-up unbiased proteomic approach^21^ with label-free peptide quantification based on our previously published data-independent-acquisition (DIA) workflow,^20^ we relatively quantified proteins based on detecting a minimum of >2 unique peptides in the PM-Mito and control groups. The mass spectrometry proteomics data have been deposited to the ProteomeXchange Consortium via the PRIDE partner repository^22^ with the dataset identifier PXD053742.

### Immunoblotting

Muscle biopsy samples were lysed in 100 μl of lysis buffer: 50 mM Tris–HCl-Applichem Biochemica A3452, pH 7.8, 150 mM NaCl (Merck), 1% SDS (Carl Roth) and supplemented with proteases/phosphatases inhibitors (Roche), using a manual glass grinder followed by sonication. Samples were centrifuged at 4 °C for 5 min at 5,000 *g*. The protein concentration of the supernatant was determined with the Pierce™ Rapid Gold BCA Protein-Assay-Kit (Thermo Fisher) according to the manufacturer’s protocol.

For immunoblotting, 10 μg of total protein extract was used for each sample analyzed. The samples were loaded on a gradient polyacrylamide gel (NuPage 4–12% Bis-Tris Protein gel, Thermo Fisher) and separated for 60 min at 150 V. Proteins were then transferred to a PVDF membrane (Invitrogen transfer stacks) using the iBlot2 dry transfer system (Thermo Fisher) according to the manufacturer’s protocol. Membranes were blocked with 5% milk powder diluted in PBS (Gibco) or TBS (Santa Cruz) with 0,1% Tween-20 (Sigma-Aldrich) for two hours and four washing steps using PBS-T. Membranes were next incubated with primary antibodies (Total OXPHOS Human WB Antibody Cocktail, Abcam, ab110411; dilution 1:800; TOM20: Recombinant Anti-TOM20 antibody, ab186735, dilution 1:2,000; GAPDH: Anti-GAPDH antibody, ab8245, dilution 1:2,000, ATP5a: Abcam, ab14748, dilution 1:2,000; CHCHD3: Proteintech, 25625-1-AP, dilution 1:5,000; ATAD3a: Abcam, ab112572, dilution: 1:1,000; SAM50: Proteintech, 28679-1-AP, dilution: 1:1,000; TIMM23: Abcam, ab230253, dilution 1:1,000; VDAC1: Abcam, ab14734, dilution: 1:1,000; STING: Cell Signaling Technologies, 13647, dilution 1:1,000; pSTING: Cell Signaling Technologies, 50907, dilution 1:1,000; CASP-1: Proteintech, 22915-1-AP, dilution 1:5,000; ACTB: Proteintech, dilution 1:5,000) at 4 °C overnight and then washed three times in 1x PBS-T. Horseradish peroxidase (HRP)-conjugated secondary goat anti-mouse antibody (Thermo Fisher Scientific) and goat anti-rabbit (Thermo Fisher Scientific) antibodies were diluted at 1:5.000 and added to the membranes for two hours. Imaging was performed after washing PVDF membranes three times in PBS-T for 10 min using enhanced chemiluminescence and a horseradish peroxidase substrate (Super-Signal West Femto, Pierce). Signals were detected using an Amersham Imager 680 machine (GE Life Sciences, USA). GAPDH was used as a loading control while mitochondrial content was normalized to the mitochondrial protein TOM20. Three control muscle biopsy specimens were used, from patients that were biopsied diagnostically for muscle pain but showed no pathological changes upon microscopic examination.

### Quantification of mitochondrial DNA copy number (mtDNA CN)

Following a previously published protocol, the relative mtDNA copy number per cell was quantified by a multiplex Taqman qPCR assay (Bio-Rad, Hercules, USA).^23^ *B2M* transcript levels were used as nuclear-encoded reference gene, and *MT-ND1* was used as mitochondrial-encoded gene. The primers used for the qPCR reaction were as follows: *B2M* Fw-CACTGAAAAAGATGAGTATGCC, Rv-AACATTCCCTGACAATCCC; *MT-ND1* Fw-ACGCCATAAAACTCTTCACCAAAG, Rv-GGGTTCATAGTAGAAGAGCGATGG. All samples were run in triplicates and replicates with a difference greater than 0.5 Ct were removed. The relative amount of mtDNA was calculated at the Ct value difference of *B2M* and *MT-ND1*, where Delta *C*_t_ (ΔC*_t_*) equals the sample C*_t_* of the mitochondrial gene (*MT-ND1*) subtracted from the sample C*_t_* of the nuclear reference gene (*B2M*). Six healthy and age-matched controls (four females and two males>50 years of age) were included for this analysis.

### Long-range PCR for mitochondrial DNA deletions

The presence of mtDNA deletions in the major arc (10kb) of the mitochondrial genome was assessed by long-range PCR. Custom-made primers were used (Fw: 5’-CCCTCTCTCCTACTCCTG-3’; Rev: 5’-CAGGTGGTCAAGTATTTATGG-3’). PCR amplification was performed on a T100 Thermal Cycler (Bio-Rad, Hercules, USA) using 1 µl of DNA and PrimeSTAR GXL DNA Polymerase (Takara Bio Europe) according to the manufacturer’s recommendations. PCR products were electrophoresed on a 0.7% agarose gel for 65 min. Three healthy and age-matched controls (two male and one female) were included in the analysis.

### Multiplexed full-length single-molecule sequencing of the mitochondrial genome by Cas9-mtDNA Oxford Nanopore sequencing

We applied a method to target, multiplex, and sequence at high coverage full-length human mitochondrial genomes as native single-molecules, utilizing the RNA-guided DNA endonuclease Cas9 (detailed methods see^24^). Combining Cas9-induced breaks that define the mtDNA beginning and end of the sequencing reads as barcodes, we achieved high demultiplexing specificity and delineation of the full-length mtDNA, regardless of the structural variant pattern. The long-read sequencing data was analyzed using a pipeline with custom-developed software, Baldur, which efficiently detects single nucleotide heteroplasmy to below 1%, physically determines the phase, and can accurately disentangle complex deletions.

### Bulk RNA sequencing

Bulk RNA sequencing (RNAseq) was performed on frozen muscle biopsy specimens as previously described.^25^ In short, RNA was extracted from fresh-frozen biopsies using TRIzol (Invitrogen) and quantified using NanoDrop. Libraries for bulk RNA sequencing were prepared using NEBNext Poly(A) mRNA Magnetic Isolation Module and Ultra II Directional RNA Library Prep Kit for Illumina (New England BioLabs, cat. #E7490 and #E7760). The input RNA and the resulting libraries were analyzed with Agilent 4200 Tapestation for quality assessment. Transcript levels are reported as log2(TMM+1) values. Histologically normal muscle biopsies were included as controls. These samples were obtained from the Johns Hopkins Neuromuscular Pathology Laboratory (n=12), the Skeletal Muscle Biobank of the University of Kentucky (n=8), and the National Institutes of Health (n=13). The normal muscle biopsies from the Johns Hopkins Neuromuscular Pathology Laboratory were obtained for clinical purposes but did not show any histological abnormality; the rest of the normal biopsies were obtained from healthy volunteers.

### Quantification of Cell-Free mtDNA from Serum

Serum samples were available from four IBM patients and two healthy controls and cryopreserved at -80°C. For DNA extraction, 100 µl of serum was centrifuged at 2000 x *g* for 5 min to remove debris and processed using the DNeasy Blood and Tissue Kit (Qiagen), with the ethanol precipitation step performed overnight at -20°C. Samples were quantified using the Qubit 1X HS dsDNA Kit (ThermoFisher), and equal volumes were loaded for mtDNA copy number qPCR as described above. Relative MT-ND1 copy number was calculated by multiplying Ct values by the calculation 2*2^-Ct^.

### Quantification and statistical analysis

Categorical variables are reported as numbers and percentages and were compared using Fisher’s exact test. Quantitative variables are reported as mean (±SD) and compared using Mann-Whitney or Kruskal-Wallis with Dunn’s multiple comparison tests. Spearman rank-order correlations were used to calculate the association’s quantitative variables. A *p*-value <0.05 was considered significant. GraphPad Prism 9.2.0 (GraphPad Software, Inc., La Jolla, CA, USA) was used for analysis.

## Data availability

The data supporting this study’s findings are available from the corresponding authors, RH and WS, upon reasonable request.

## Results

### Clinical and demographic data of the patient cohorts

Our study included skeletal muscle biopsy specimens from 27 patients with PM-Mito (22 female, five male) and 27 patients with typical IBM (13 female, 14 male), meeting the ENMC criteria for clinicopathologically defined IBM.^15^ We previously reported the clinical features of these cohorts in detail.^2^ Significantly, the two patient groups did not differ in age at biopsy (PM-Mito 66 ± 12 years, IBM 69 ± 9.2 years, *p*=0.25). While all patients with IBM presented with the pathognomonic clinical pattern, PM-Mito patients showed a spectrum of clinical symptoms. The most common clinical presentations at the time of muscle biopsy for PM-Mito patients were proximal muscle weakness (bilateral quadriceps femoris muscle weakness) and other incomplete IBM-like patterns not meeting ENMC criteria (e.g., isolated knee extensor weakness, long finger and wrist flexor weakness). Creatine kinase (CK) levels were moderately elevated (up to 10x the upper limit of normal (≥174 U/l)) in both groups, again without statistically significant differences (data not shown). As previously reported,^26^ all late IBM and 16/27 PM-Mito patients have received immunosuppressive treatment, which has not led to sustained clinical improvement in any of the patients.

### Functional, structural, and molecular alterations of mitochondria

#### COX-negative/SDH-positive muscle fibers were present in both PM-Mito and IBM muscle

Twenty-seven muscle biopsy specimens derived from PM-Mito patients were available for comprehensive histological analysis, and 27 randomly selected specimens showing classic IBM features were studied. All muscle biopsy samples showed a combination of mild to moderate fibre-size variation with endomysial lymphomonocytic infiltrates, mildly increased numbers of internalized myonuclei, and intense sarcolemmal staining by MHC class I (Fig. 1). Focal MHC class II positivity was observed, particularly in areas with CD8+ T-cell infiltrates. As published previously, vacuoles and p62-positive intracellular aggregates were absent in all PM-Mito cases,^2^ as these would classify these cases as IBM according to the ENMC criteria.

**Figure 1.**
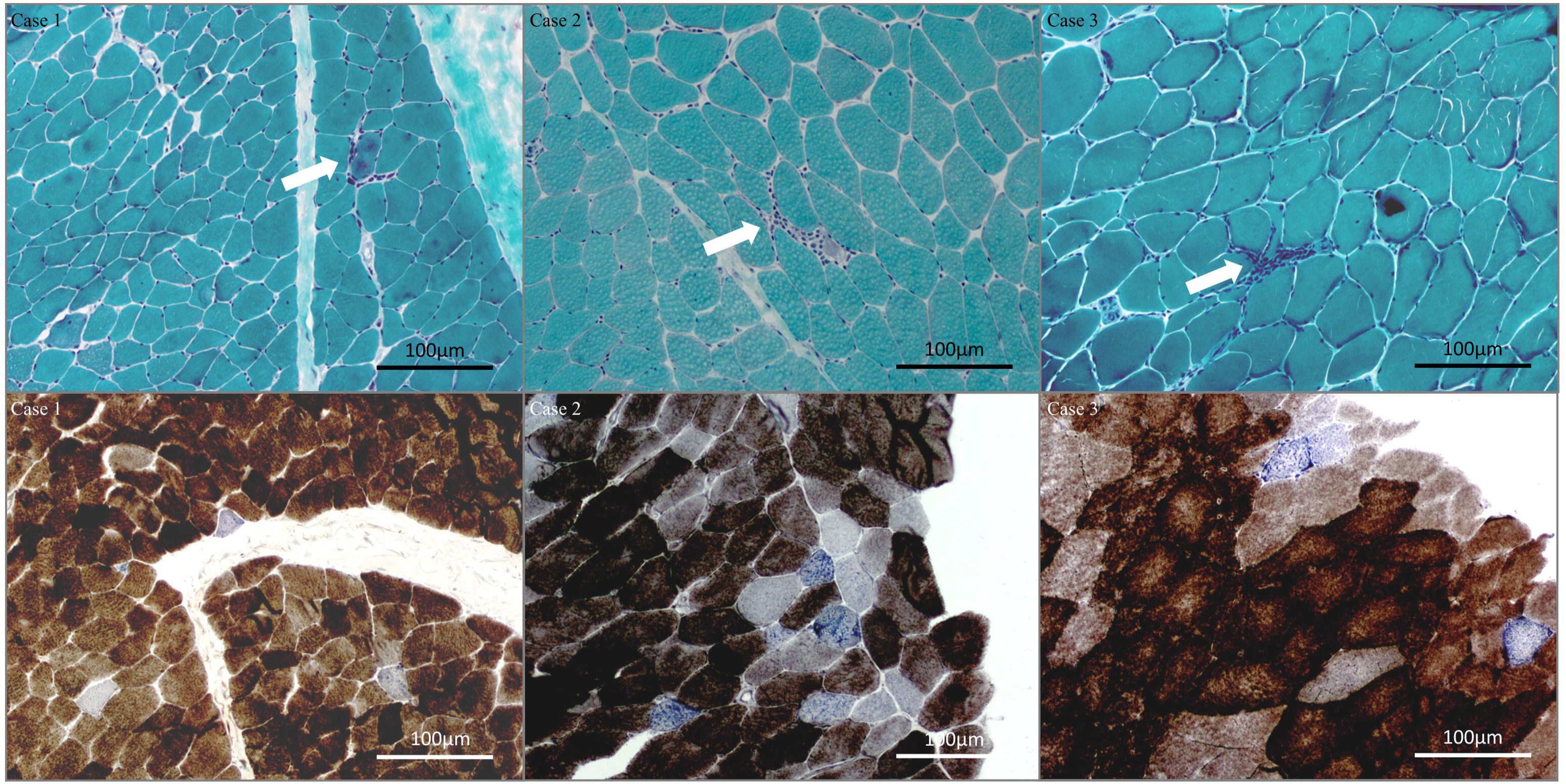
Inflammation and mitochondrial dysfunction in three cases of PM-Mito. **Cases 1-3** illustrate muscle biopsy specimens with few inflammatory infiltrates (arrows) but pronounced mitochondrial abnormalities, evident as COX-deficient myofibers. Top row: Gömöri-trichrome stain. Bottom row: COX-SDH histochemistry.

PM-Mito skeletal muscle biopsy specimens revealed small clusters of infiltrates (Fig. 1, top row) of T-cells within the endomysium, often encircling individual myofibers or small groups of myofibers with interspersed macrophages. All PM-Mito muscle biopsy specimens showed an increased amount of COX-negative (SDH-positive) muscle fibers for the patients’ age according to established criteria (i.e., >3% of muscle fibers).^27^ Quantifying COX-negative muscle fibres revealed no statistical difference in fibres exhibiting signs of mitochondrial dysfunction between PM-Mito cases (25.6 ± 21.4) and typical IBM cases (34 ± 23.6, *p*=0.19). Mitochondrial abnormalities were evident throughout all stages of the disease, including very mild cases with few inflammatory infiltrates. Three representative PM-Mito cases showing only mild myopathic alterations, without evident fatty-fibrotic tissue remodeling and few infiltrates but pronounced mitochondrial abnormalities in COX/SDH stains, illustrated in Fig. 1.

#### Distinct ultrastructural mitochondrial abnormalities in PM-Mito and IBM

TEM was performed on ten randomly selected PM-Mito cases and ten IBM cases. Overall, in TEM studies, all muscle biopsy specimens showed mitochondrial abnormalities with quantitative rather than qualitative differences between the individual patients. Non-specific signs such as swollen, elongated, and dysmorphic mitochondria and subsarcolemmal accumulation were present in all PM-Mito and IBM cases. We also observed discontinuity of the mitochondrial membranes, accompanied by leakage of mitochondrial content into the sarcoplasm (Fig. 2A). Other features included paracrystalline inclusions (PCI, Fig. 2B) type 1 and 2, concentric cristae (CC, Fig. 2C), and giant mitochondria with densely packed cristae membranes (GM,^28^ Fig. 2D). We quantified these distinct ultrastructural abnormalities in PM-Mito and IBM cases at 12.000- to 20.000-fold magnification. PCI were detected in 8.7% of PM-Mito and IBM in 12.6% of mitochondria. CC was found in 10,8% and 11,8%, and GM in 0.6% and 12.6% of mitochondria, respectively. Mitophagy was occasionally seen but not a prominent ultrastructural finding. In summary, ultrastructural abnormalities of mitochondria were detected throughout all stages of the disease, including cases showing very mild signs of inflammation. However, giant mitochondria with densely packed cristae were more abundant in typical IBM cases.

**Figure 2.**
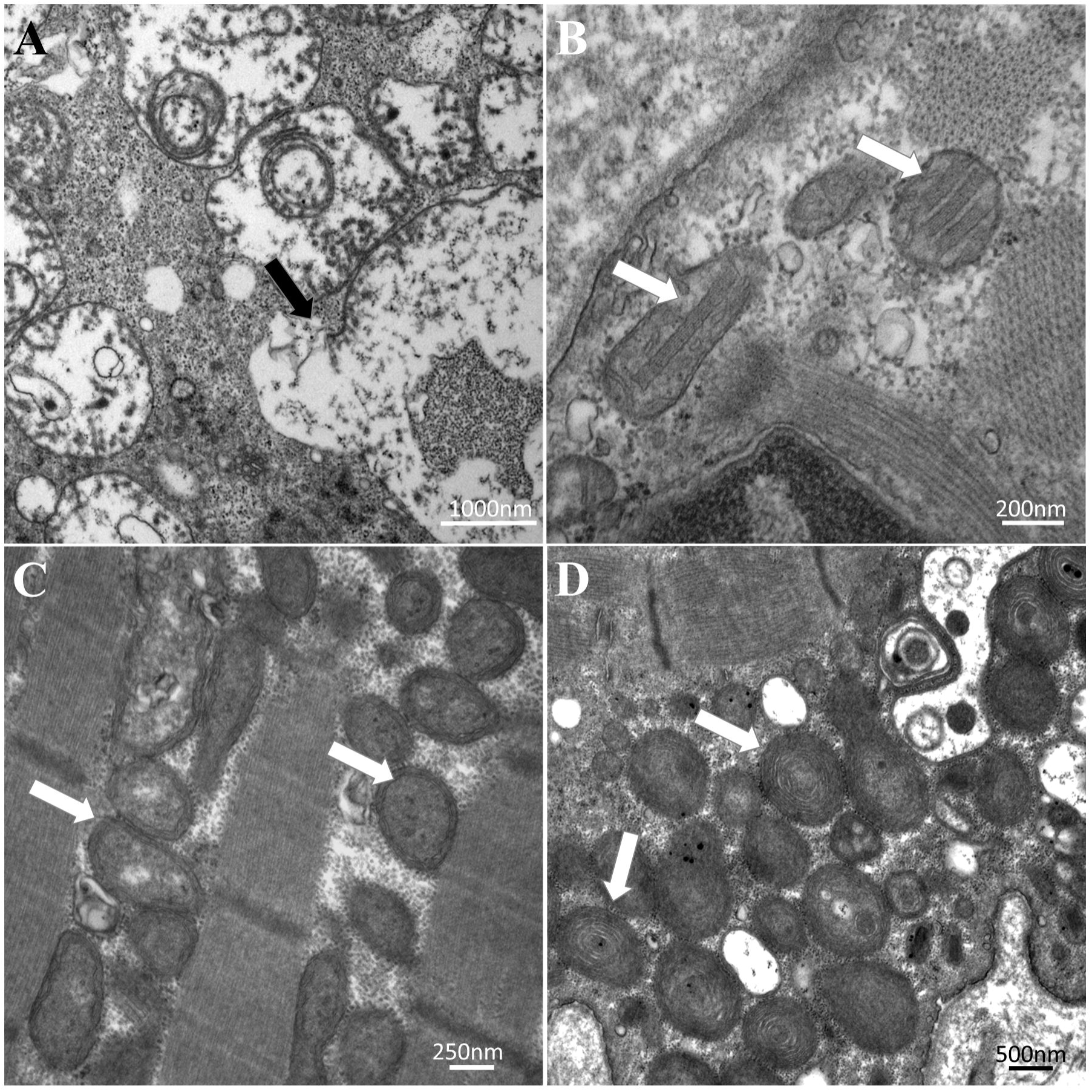
Ultrastructural features of abnormal mitochondria in IBM-SD. Representative images of mitochondrial pathology in PM-Mito **(A-C)** and IBM **(D)**. **(A)** Disruption of mitochondrial membranes and leakage of mitochondrial content (arrows) **(B)** Paracrystalline inclusions (arrows) **(C)** Circular mitochondrial membranes (with lack of cristae) (arrows) **(D)** Mitochondria with densely packed concentric cristae (arrows).

#### Multiple mtDNA deletions and depletion are early signs of IBM-spectrum disease

PCR-based mtDNA copy number measurements were performed on ten PM-Mito, eight IBM, and 13 control muscle biopsy specimens. The level of mtDNA showed a statistically significant reduction in PM-Mito (7.1 ± 0.91) and typical IBM patients (6.57 ± 0.88) compared to healthy, old-age control specimens (8.2 ± 0.61, Fig. 3A). Of note, there was no difference between the PM-Mito and IBM groups (*p*=0.33), however mtDNA copy numbers were significantly different in both PM-Mito (*p*=0.0077) and IBM (*p*=0.0002) than controls, indicating that the mtDNA defect is maintained throughout all stages of the disease.

**Figure 3.**
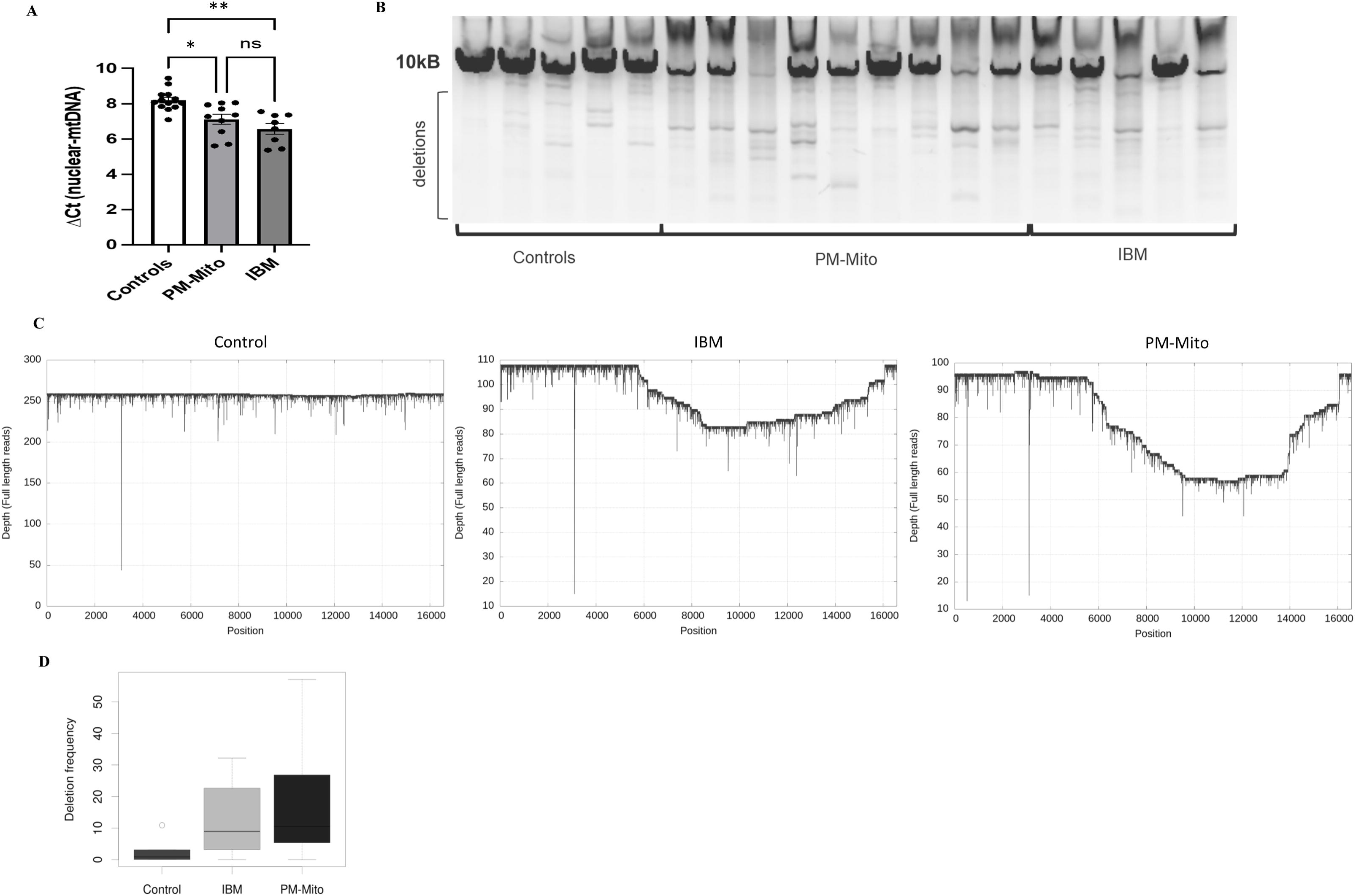
Alterations of mtDNA in PM-Mito and typical IBM. (**A**) qPCR based mtDNA copy number assessment showing a reduction in copy numbers in IBM-SD patients, but no significant difference between PM-Mito and IBM patients **(B)** Long-range PCR of the mtDNA major arc showing multiple, age-inappropriate mtDNA deletions in PM-Mito and IBM muscle biopsy specimens, compared to age-matched controls. **(C)** Multiplexed full-length single-molecule sequencing of mtDNA detected several medium and large-scale deletions with similar deletion patterns in PM-Mito and IBM. The multiple mtDNA deletions were detected in heteroplasmy rates between ∼1% to >50%, and often covered regions from 8-12kb, but deleted regions spanned 3-16kb. The amount of mtDNA deletions was significantly higher in patients compared to age-matched controls, and the heteroplasmy rate was slightly higher in PM-Mito than in late IBM. \**p*=0.0077 \*\**p*=0.0002

To further investigate a possible link between mitochondrial dysfunction and deletions of the mtDNA, we performed a long-range PCR of the major arc, where most mtDNA deletions are known to be located.^29^ Nine PM-Mito, five typical IBM, and five control cases were studied using this technique. As mtDNA deletions are known to accumulate in aging skeletal muscle, controls representing a similar age range (50 to 83 years) were included. Indeed, we detected a similarly high number of multiple mtDNA deletions in PM-Mito and IBM (Fig. 3B), suggesting that mtDNA deletions arise early in the pathophysiology of the disease.

#### Multiplexed full-length single-molecule sequencing of the mitochondrial genome

We detected a large number of multiple mtDNA deletions of different sizes in both PM-Mito and IBM samples (Fig. 3C). All samples showed deletions with a range of sizes and heteroplasmy rates between ∼1% and >50% (Median ∼11%). Deletions tend to cover regions from 8-12kb, but deleted regions often span large regions from 3-16kb. The number of single nucleotide mutations in the mtDNA was less prominent in the patient samples than mtDNA deletions.

The number of mtDNA deletions was significantly higher in IBM-SD (IBM and PM-Mito) patient samples than in controls (*p*=0.025) but not significantly different between PM-Mito and IBM patients (*p*=0.82; Fig. 3C, D). The heteroplasmy rate was slightly higher in PM-Mito than in IBM, but not reaching statistical significance. These results confirm data obtained by long-range PCR analysis that the mtDNA alterations are early signs of the pathology.

#### Unbiased proteomic profiling and bulk RNAseq

We performed unbiased proteomic profiling of 13 PM-Mito muscle biopsy specimens. We detected 3787 proteins using this approach, of which 33 were statistically significantly downregulated, and 20 were upregulated in PM-Mito patient samples vs. controls (Fig. 4A). Interestingly, 11/33 downregulated proteins were linked to mitochondrial function, including complexes I and III of the respiratory chain, and two out of the ten most downregulated proteins were related to the inner mitochondrial membrane (SMDT1, TIMM21). The most upregulated protein, with a 6.1-fold upregulation, was TMEM14C (Transmembrane protein 14C), a protein located in the inner mitochondrial membrane and involved in various mitochondrial processes, including heme biosynthesis.

**Figure 4.**
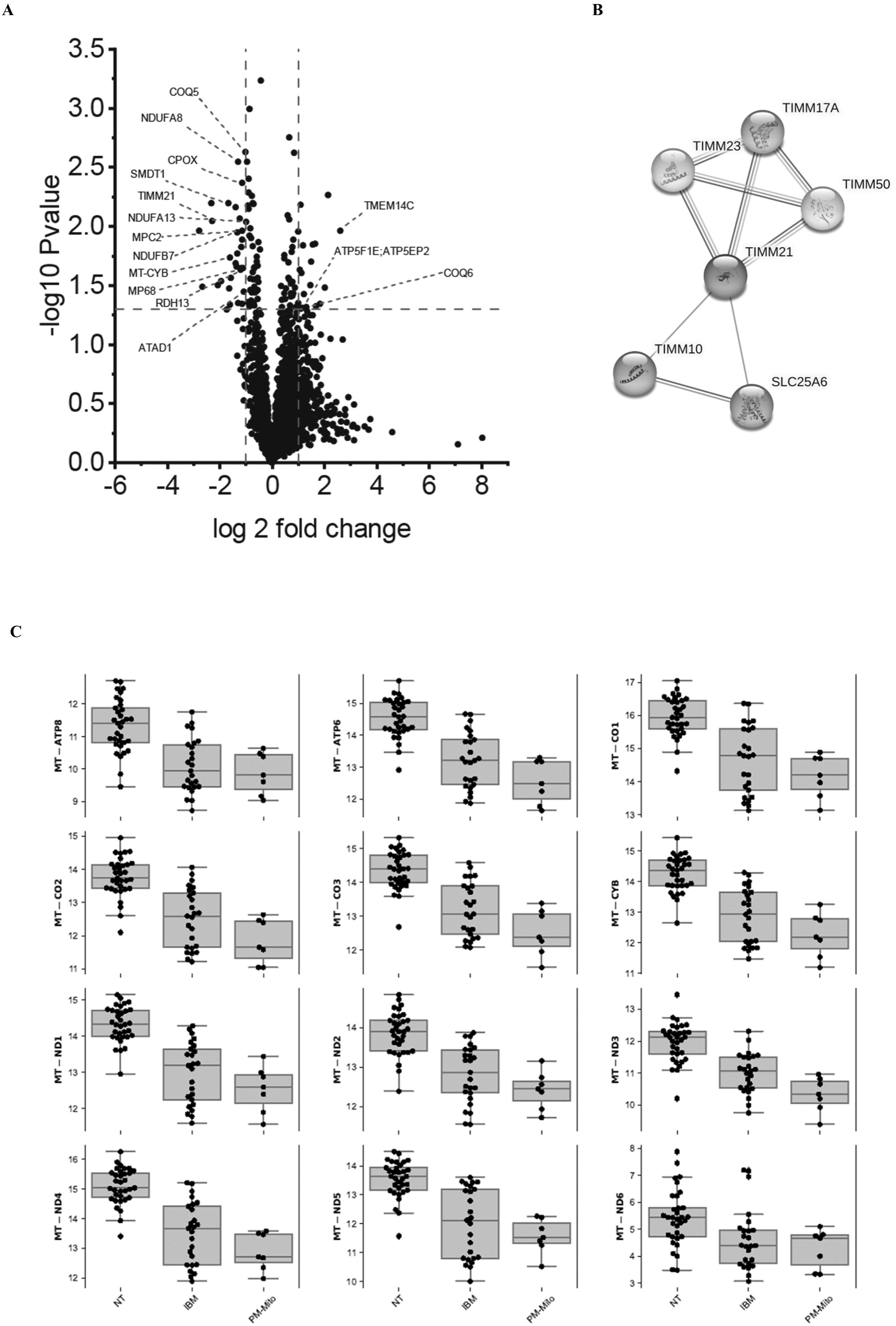
Unbiased proteomic profiling and bulk RNAseq of PM-Mito and typical IBM muscle biopsy specimens. (**A**) Volcano plot illustrating dysregulated mitochondrial proteins in PM-Mito, showing a profound dysregulation of proteins associated with complexes I and III, and inner mitochondrial membrane proteins (**B**) STRING illustration of the TIMM21 interaction network including TIMM23, both being part of the inner mitochondrial membrane **(C)** Downregulation of mtDNA encoded mitochondrial genes, affecting all mtDNA encoded respiratory chain subunits in PM-Mito and IBM. Graphs represent log-scaled normalized expression (log2(TMM+1)) levels of differentially expressed mRNA in PM-Mito, IBM vs. controls (NT).

Gene ontology (GO) term analysis indicated that respiratory chain/mitochondrial function and cytoskeletal remodeling were the most significantly downregulated biological processes. On the other hand, chromatin silencing appeared as the most upregulated biological process in this analysis, hinting toward epigenetic changes.

Bulk RNAseq of nine IBM and seven PM-Mito muscle biopsy specimens showed significant downregulation of both nuclear- and mtDNA-encoded mitochondrial transcripts in PM-Mito and IBM, thus confirming the downregulation observed on the protein level. There were no apparent differences in expression levels between the patient groups (Fig. 4C; Supplementary Fig. 1).

#### Immunoblotting of mitochondrial membrane-associated proteins

To study the molecular underpinnings of morphological and ultrastructural abnormalities of mitochondria, we immunoblotted mitochondrial membrane-associated proteins in three PM-Mito and three IBM patient samples compared to three control samples. There was a marked decrease in key inner mitochondrial membrane proteins, including ATP5A, CHCHD3/MIC19, VDAC1, and TIMM23 (Fig. 5A and 5B). Unfortunately, in one PM-Mito sample, successful immunoblotting was achieved only for CHCHD3 and ß-Actin. Due to insufficient material, it was not possible to repeat the immunoblotting on this sample.

**Figure 5.**
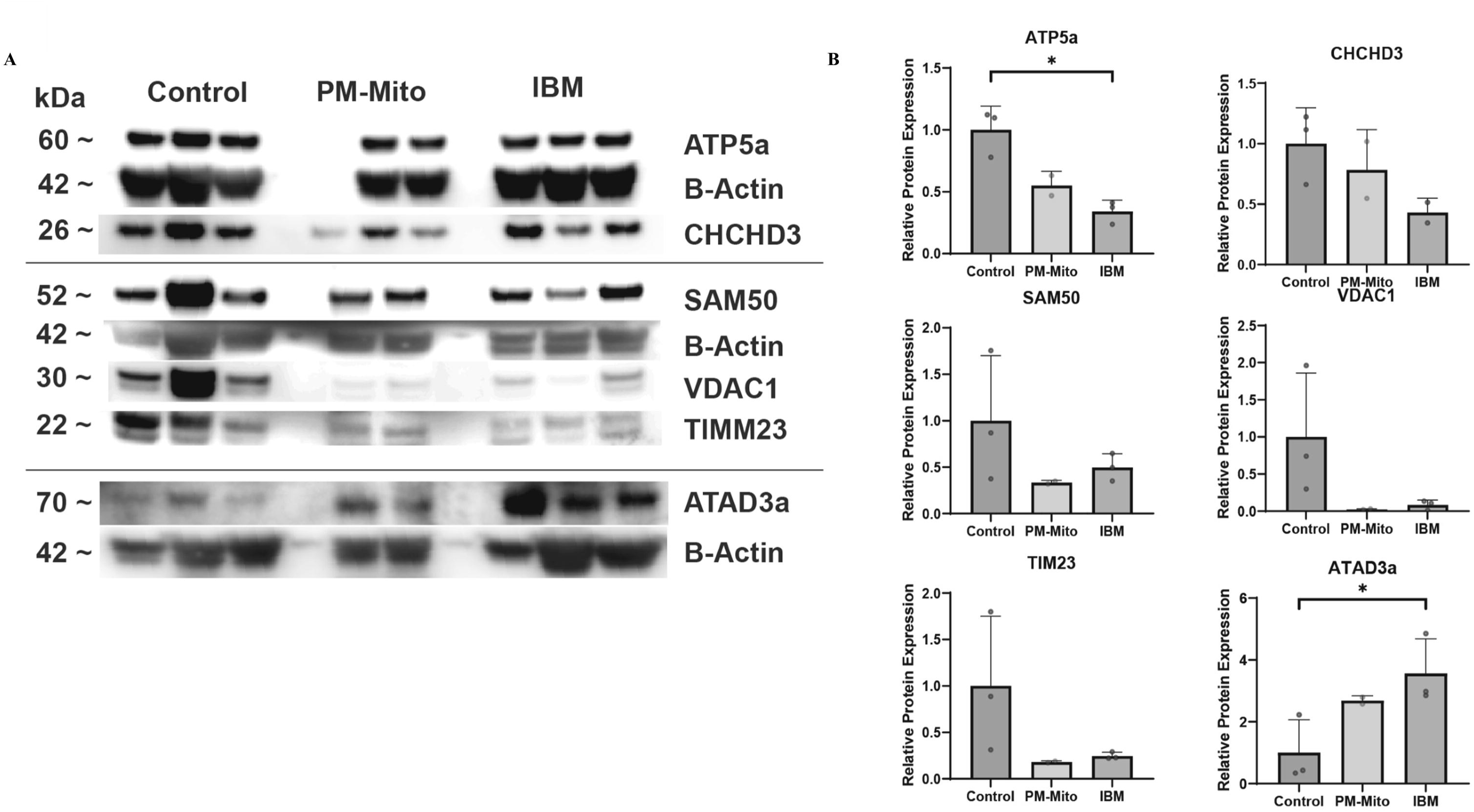
Structural and functional alterations of mitochondria in PM-Mito. (**A**) Representative images of immunoblotting for mitochondrial membrane proteins illustrating dysregulation of mitochondrial membrane proteins in PM-Mito, compared to typical IBM. **(B)** Quantification of protein expression normalised to β-Actin and relativized to controls average. \**p*<0.05

Recent work has shown that a functional complex forms between mitochondrial proteins SAM50, ATAD3A, and MIC19/MICOS to maintain membrane and mtDNA stability.^30^ All the proteins play key roles in the organization and maintenance of the mitochondrial membranes, including the cristae structure. This molecular interaction is also illustrated by STRING analysis of differentially downregulated proteins, with TIMM21 showing a close interaction with TIMM23 (Fig. 4B).

### Inflammation and mitochondrial dysfunction in PM-Mito and IBM

#### Activation of canonical pro-inflammatory pathways

It has previously been described that mitochondrial damage can lead to the intracellular leakage of mtDNA, which may activate the pro-inflammatory cGAS/STING pathway.^31^ Therefore, after confirming the presence of widespread mitochondrial abnormalities in PM-Mito, we studied the expression levels of proteins linked to the cGAS/STING pathway by immunoblotting. We detected an upregulation of cGAS downstream targets pSTING, total STING, and pro and active forms of Caspase-1 in both PM-Mito and IBM but not in control samples (Fig. 6A, 6B). Caspase-1 is closely linked to the cGAS/STING-mediated inflammasome assembly.^31^

**Figure 6.**
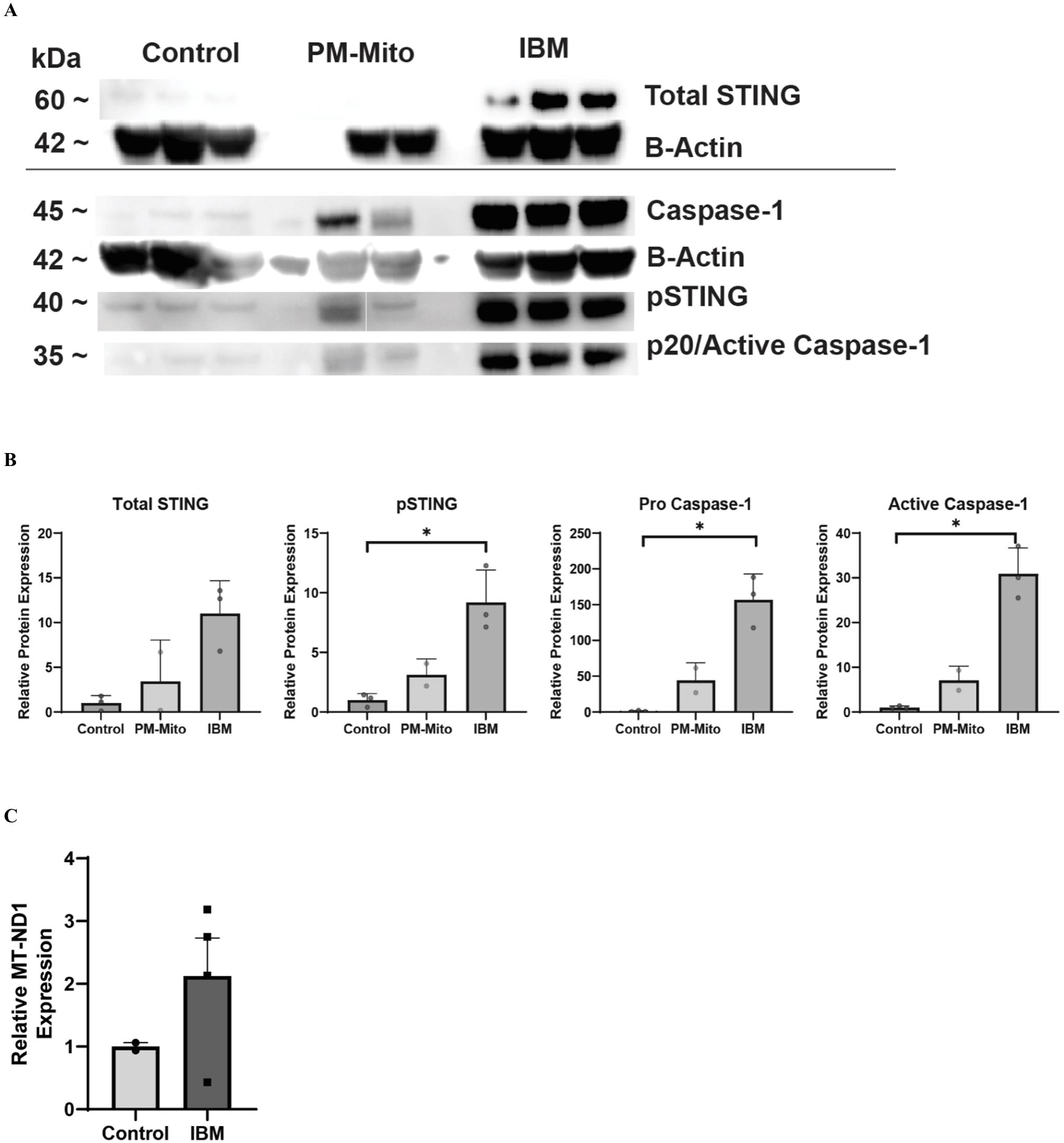
Mitochondria-associated inflammation in PM-Mito and typical IBM patients. (**A**) Representative images of immunoblotting for cGAS/STING-associated proteins and Caspase-1 (**B**) Quantification of protein expression normalised to β-Actin and relativized to controls average **(C)** Cell-free mtDNA quantification from the serum of IBM patients showed elevated levels of circulating mtDNA in the patient group, indicating a leakage of mtDNA from muscle cells into the blood. \**p*<0.05

Bulk RNAseq confirmed an upregulation of the cGAS/STING pathway in PM-Mito and typical IBM compared to controls at gene expression level. Interestingly, cGAS/STING activation was more pronounced in PM-Mito samples at the RNA level (Supplementary Fig. 2). Further correlation analysis showed that cGAS/STING-associated transcript levels showed positive correlation with immune cell markers (e.g., CD8, CD68), and negative correlation with markers of muscle regeneration and structure (e.g., MYH3, ACTA1; Supplementary Fig. 3).

To confirm if this activation in DNA sensing pathways might be triggered by the leakage of mtDNA, patient serum samples were analyzed for cell-free mtDNA (cfmtDNA) to assess the extracellular release of mtDNA. CfmtDNA has been shown to act as a potent pro-inflammatory trigger.^32^ IBM samples showed more circulating cfDNA than controls, doubling the circulating mtDNA levels (Fig. 6C). This indicated that mtDNA release from the mitochondria is prominent in IBM and may be a potential inflammatory trigger.

#### Inflammatory infiltrates and the extent of mitochondrial pathology

To study whether the amounts of infiltrating immune cells (CD8+ T-cells, CD68+ macrophages, CD45+ leukocytes) and COX-SDH-positive cells correlated in the PM-Mito and IBM cohorts, we performed Pearson correlation analysis. This revealed no correlation between T-cell-mediated inflammation and the extent of mitochondrial alterations (i.e., COX-SDH-positive fibers) in the PM-Mito (*p*=0.6) and typical IBM (*p*=0.98) cohorts. Furthermore, neither the number of CD68+ macrophages (PM-Mito *p*=0.93; typical IBM *p*=0.95) nor the total CD45+ leukocyte count (PM-Mito *p*=0.61; typical IBM *p*=0.90) correlated with COX-SDH abnormalities in the patient groups. There was no spatial association between COX-SDH abnormalities and the distribution of inflammatory infiltrates.

## Discussion

This study focused on mitochondrial alterations as an early feature of IBM pathophysiology. To study this, we investigated PM-Mito as an early form of IBM. We observed quantitatively and qualitatively similar mitochondrial abnormalities at the histological, ultrastructural, and molecular levels in both PM-Mito and typical IBM. The number of COX-negative fibers were similar, and higher than expected by the age of the patients in both PM-Mito and IBM. The extent of mitochondrial damage did not correlate with the amount of inflammation, suggesting no causative effect. Abnormalities such as disorganized and circular mitochondrial membranes, along with enlarged and dysmorphic mitochondria, were prevalent in both PM-Mito and IBM patients. Interestingly, ultrastructural abnormalities also included disturbed cristae architecture and discontinuity of the mitochondrial membranes, possibly allowing a leakage of mitochondrial content into the sarcoplasm. Multiple large-scale mtDNA deletions and reduced copy numbers were evident in PM-Mito, compared to healthy age-matched controls. Notably, there was no significant difference in these mitochondrial DNA alterations between the PM-Mito and IBM groups. Additionally, structural proteins of the inner mitochondrial membrane and proteins involved in membrane and mtDNA quality control (SAM50, ATAD3A, MIC19/MICOS) were differentially regulated in PM-Mito and IBM samples. Immunoblotting confirmed dysregulation of proteins related to mitochondrial ultrastructure, thus validating data on unbiased proteomic and transcriptomic profiling. We also detected the activation of the canonical cGAS/STING pathway throughout all stages of IBM-SD (i.e, in both PM-Mito and IBM), highlighting that the intracellular release of mitochondrial DNA, followed by activation of the innate immune response further contributes to the pathomechanism.

The data reported here corroborate our earlier findings,^2^ indicating that mitochondrial dysfunction is an early feature of IBM spectrum disease. Therefore, mitochondrial dysfunction should be regarded as closely linked to the disease pathogenesis rather than simply a consequence of prolonged exposure to inflammation. Along this line, our data clearly show that inflammatory infiltrates do not correlate to the extent of mitochondrial damage at the morphological level, and the mitochondrial dysfunction is not simply a secondary effect of inflammation.

The role of mitochondria in the pathogenesis of IBM is undoubtedly both significant and complex. In general, mitochondrial abnormalities are not a typical feature of idiopathic inflammatory myopathies, with the exception perhaps of dermatomyositis (DM) showing perifascicular COX-SDH abnormalities.^33,34^ However, the molecular mechanisms underlying mitochondrial dysfunction in DM differ from those observed in IBM, as single point mutations and depletion of the mtDNA in DM^35^ have been restricted to focal areas with inflammatory infiltrates.^36^ In contrast, IBM muscle is characterized by large-scale deletions in the major arc of mtDNA molecules^37^ and lower mtDNA copy numbers than healthy subjects of similar age groups.^38^

We could reproduce these findings in both cohorts of PM-Mito and IBM patients. At the protein level, combined unbiased proteomic and immunoblotting both point toward an involvement of the inner mitochondrial membrane, which is underlined by STRING analysis showing that the most downregulated inner mitochondrial membrane protein, TIMM21, is a close interaction partner of TIMM23, which we also found downregulated in the immunoblot. The disruption of this complex protein network possibly results in disturbed ultrastructure and defective mtDNA replication. It has been previously shown that the morphology of mitochondria in IBM myofibers is altered, with features such as shortened and enlarged cristae, membrane disruption, and a reduced mitochondrial length-to-width ratio.^39^ We observed giant mitochondria with highly abnormal cristae, which were almost exclusively detected in IBM, but not in PM-Mito, suggesting that these ultrastructural abnormalities develop at a later stage. This peculiar ultrastructural abnormality has already been described in the context of IBM.^28,40^ It is unclear whether this is linked to disease duration, but patients with mitochondrial myopathy caused by *m.8344A>G* mutations show similar ultrastructural abnormalities.^41^ Our ultrastructural data clearly showed similar patterns in PM-Mito and IBM, underlining the functional impact of the dysregulation of (inner) mitochondrial membrane proteins.

Another relevant ultrastructural feature observed in this study was the disruption of mitochondrial membranes and leakage of mitochondrial content into the sarcoplasm. The release of mtDNA and mitochondrial content into the cytosol has been described to act as mitochondrial-derived damage-associated molecular pattern (DAMP),^32^ triggering inflammatory cascades such as the cGAS/STING pathway.^31,42^ DAMPs are recognized as foreign by the immune system’s pattern recognition receptors and this triggers the activation of pro-inflammatory pathways such as toll-like receptor (TLR) signaling.^43^ For example, the binding of oxidized cardiolipins to TLR4 in the cytosol can initiate NF-κB signaling, leading to increased myostatin expression, which has been detected in IBM muscle.^44^ Thus, it is possible that the profound mitochondrial abnormalities observed in PM-Mito could trigger canonical inflammatory pathways, further contributing to the disease progression. Testing this hypothesis, we observed an upregulation of the cGAS/STING pathway at both RNA and protein levels in our PM-Mito and IBM cohorts. This finding is particularly relevant, as it indicates a significant role of mitochondria-associated inflammation in the early pathophysiology of IBM-SD, i.e., before fatty-fibrotic tissue remodeling and massive lymphocytic infiltration are observed. Interestingly, it has been shown that IBM is characterized by a distinct interferon signature at the gene expression level, with a predominance of type II interferons.^45^ The downstream effects of STING, a potent transcriptional activator of interferon-stimulated genes could explain this signature.^46^ This activation of pro-inflammatory cascades was also evident in our RNAseq data. On the morphological level, the prominent upregulation of Major Histocompatibility Complex (MHC) class I in PM-Mito also clearly indicates a role for interferon type I in the pathogenesis of the disease. In this context, the release of IFN-II (i.e., IFN-gamma), which has been well described in IBM, may be explained by downstream effects of IFN-I and persistence of a pro-inflammatory milieu, leading to the invasion of IFN-gamma releasing immune cells (e.g., CD4+ T-cells, macrophages). However, this interplay warrants further investigation. Our data indicate that these immune phenomena might be secondary to mitochondrial dysfunction and associated inflammation in the IBM cascade. This, in turn, could suggest that IBM can be viewed as a primarily degenerative disease with peculiar (auto-)inflammatory features. Indeed, recently published data from mouse xenograft models pointed towards IBM being a primarily degenerative disease of skeletal muscle,^47^ in which inflammation may be a bystander or a secondary phenomenon. However, the initial triggers of mitochondrial dysfunction in IBM remain unclear.

We detected large-scale deletions of mtDNA in PM-Mito patients, raising the possibility that mtDNA damage could be a primary cause of mitochondrial dysfunction in IBM-SD. Several explanations for these mtDNA deletions in the context of IBM have been discussed: errors in DNA replication, ROS accumulation, impaired DNA repair systems, and defective autophagy (mitophagy).^48^ Moreover, the reduced mtDNA copy numbers point towards an mtDNA maintenance defect. Notably, single nucleotide polymorphisms (SNPs) in key genes involved in mtDNA replication and maintenance, such as the DNA helicase Twinkle, DNA polymerase γ (*POLG*), and ribonucleotide-diphosphate reductase subunit M2B (*RRM2B*), have been identified in IBM patients.^49^ However, no significant differences in variant prevalence in *POLG* were found between the IBM groups or between IBM patients and the control population. Thus, the pathogenic role of these SNPs has not been convincingly demonstrated.^49^ We hypothesize that the multiple mtDNA deletions and depletion may be related to the severe abnormalities of mitochondrial cristae, where mtDNA replication occurs. However, understanding the exact mechanism of these events requires further studies.

Another interesting hypothesis, derived from studies performed in the context of amyotrophic lateral sclerosis (ALS), links mitochondrial damage and the release of mtDNA to the deposition of TDP-43 in the cytosol and in mitochondria.^50^ Indeed, TDP-43 mislocalization and sarcoplasmic deposition are characteristic, but not specific, features observed in IBM muscle.^51^ Others and we recently demonstrated the presence of TDP-43-associated cryptic exons in IBM muscle,^2,47^ pointing towards a functional impact of TDP-43 pathology. However, we showed that the quantity of cryptic exon inclusion was relatively low in PM-Mito compared to typical IBM muscle. This argues against a functional link between early mitochondrial dysfunction and TDP-43 deposition.

Lastly, from a clinical perspective, patients with PM-Mito present a spectrum of clinical symptoms, including non-specific complaints such as myalgia. Defining the onset of the disease in this cohort is challenging, as IBM-SD may begin insidiously with vague, non-specific symptoms. Therefore, previous and present studies may have missed even earlier forms of the disease, and the progression of this diverse phenotype into the characteristic clinical pattern of IBM remains unclear. Recently published studies on muscle MRI patterns in PM-Mito have yielded conflicting results, showing muscle involvement patterns similar to IBM in one^52^ and partly different patterns in another study.^53^ However, the data reported here again illustrate the considerable molecular overlap between PM-Mito and IBM, underlining the importance of considering IBM as a disease spectrum with early and late stages.

In conclusion, this study established mitochondrial abnormalities as an early and characteristic finding in IBM-SD. PM-Mito and typical IBM did not differ significantly regarding mitochondrial morphology, ultrastructure, and molecular characteristics of mitochondrial damage. Importantly, our data indicate that inflammation does not trigger mitochondrial dysfunction in a time-dependent manner but points towards mitochondrial dysfunction being an early event in disease pathogenesis. Considering ongoing IBM clinical trials with immunosuppressants, this finding may have important implications for clinical practice.

**Table 1.**
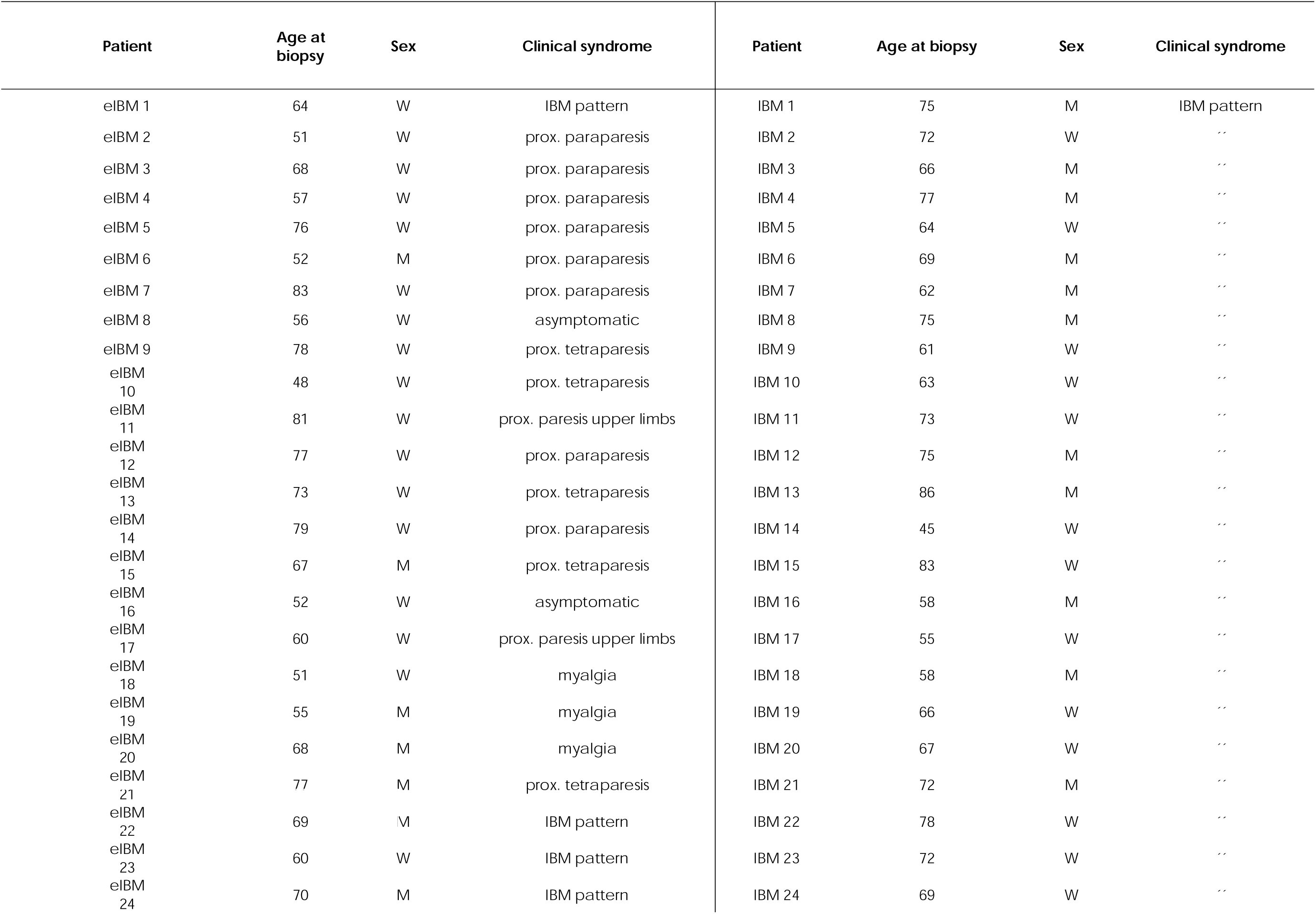
Demographic and clinical data.

## Funding

We thank the Deutsche Gesellschaft für Muskelkranke (DGM) e.V. for funding this study. AH acknowledges the support by the “Ministerium für Kultur und Wissenschaft des Landes Nordrhein-Westfalen” the “Regierenden Bürgermeister von Berlin-Senatskanzlei Wissenschaft und Forschung” and the “Bundesministerium für Bildung und Forschung.” The European Regional Development Fund (ERDF) financed parts of this study in the framework of the NME-GPS project (grant to AR). R.H. is supported by the Wellcome Discovery Award (226653/Z/22/Z), the Medical Research Council (UK) (MR/V009346/1), the Addenbrookes Charitable Trust (G100142), the Evelyn Trust, the Stoneygate Trust, the Lily Foundation, Ataxia UK, Action for AT, the Muscular Dystrophy UK, the LifeArc Centre to Treat Mitochondrial Diseases (LAC-TreatMito) and the UKRI/Horizon Europe Guarantee MSCA Doctoral Network Programme (Project 101120256: MMM). She is also supported by an MRC strategic award to establish an International Centre for Genomic Medicine in Neuromuscular Diseases (ICGNMD) MR/S005021/1 and by the NIHR Cambridge Biomedical Research Centre (BRC-1215-20014). The views expressed are those of the authors and not necessarily those of the NIHR or the Department of Health and Social Care. CNAG: This research has received funding from the European Union’s Horizon 2020 research and innovation programme under grant agreement No. 824110 – EASI-Genomics.

## Competing interests

Katrin Hahn received financial reimbursement for consulting, advisory board activities, speaker fees and/or contributions to congresses and travel support to attend scientific meetings by Akcea Therapeuticals Inc., Alnylam Pharmaceuticals Inc., Amicus, AstraZeneca, GSK, Hormosan, Takeda Pharmaceutical Inc., Pfizer Pharmaceuticals Inc., Swedish Orphan Biovitrum Inc. and ViiV Healthcare GmbH. K.H. further received research funding by the foundation Charité (BIH clinical fellow), Alnylam Pharmaceuticals Inc., and Pfizer Pharmaceuticals. None of this is related to the present work.

## Supplementary material

Supplementary material is available at *Brain* online.

**Supplementary Figure 1** Graphs represent log-scaled normalized expression (log2(TMM+1)) levels of differentially expressed mRNA of nDNA and mtDNA encoded mitochondrial RNA in PM-Mito, IBM vs. controls (NT).

**Supplementary Figure 2** Graphs represent log-scaled normalized expression (log2(TMM+1)) levels of differentially expressed mRNA related to the cGAS/STING pathway in PM-Mito, IBM vs. controls (NT).

**Supplementary Figure 3** Correlation of the most differentially overexpressed genes in patients with PM-Mito, and markers of muscle differentiation (*NCAM1*, *MYOG*, *PAX7*, *MYH3*, *MYH8*) and structural mature muscle proteins (*ACTA1*, *MYH1*, *MYH2*).

## References

1. Blume G, Pestronk A, Frank B, Johns DR. Polymyositis with cytochrome oxidase negative muscle fibres. Early quadriceps weakness and poor response to immunosuppressive therapy. Brain. 1997;120(1):39–45.

2. Kleefeld F, Uruha A, Schänzer A, et al. Morphological and Molecular Patterns of Polymyositis With Mitochondrial Pathology and Inclusion Body Myositis. Neurology. Oct 4 2022;

3. Santorelli FM, Sciacco M, Tanji K, et al. Multiple mitochondrial DNA deletions in sporadic inclusion body myositis: a study of 56 patients. Annals of neurology. Jun 1996;39(6):789–95.

4. Horvath R, Fu K, Johns T, Genge A, Karpati G, Shoubridge EA. Characterization of the mitochondrial DNA abnormalities in the skeletal muscle of patients with inclusion body myositis. J Neuropathol Exp Neurol. May 1998;57(5):396–403.

5. Hedberg-Oldfors C, Lindgren U, Basu S, et al. Mitochondrial DNA variants in inclusion body myositis characterized by deep sequencing. Brain Pathol. May 2021;31(3):e12931.

6. Zhou M, Cheng X, Zhu W, et al. Activation of cGAS-STING pathway – A possible cause of myofiber atrophy/necrosis in dermatomyositis and immune-mediated necrotizing myopathy. Journal of Clinical Laboratory Analysis. 2022;36(10):e24631.

7. Wilkinson MGL, Moulding D, McDonnell TCR, et al. Role of CD14+ monocyte-derived oxidised mitochondrial DNA in the inflammatory interferon type 1 signature in juvenile dermatomyositis. Annals of the Rheumatic Diseases. 2023;82(5):658.

8. Skopelja-Gardner S, An J, Elkon KB. Role of the cGAS-STING pathway in systemic and organ-specific diseases. Nat Rev Nephrol. Sep 2022;18(9):558–572.

9. Huntley ML, Gao J, Termsarasab P, et al. Association between TDP-43 and mitochondria in inclusion body myositis. Laboratory Investigation. 2019/07/01/ 2019;99(7):1041–1048.

10. Askanas V, Engel WK. Sporadic inclusion-body myositis: Conformational multifactorial ageing-related degenerative muscle disease associated with proteasomal and lysosomal inhibition, endoplasmic reticulum stress, and accumulation of amyloid-β42 oligomers and phosphorylated tau. La Presse Médicale. 2011/04/01/ 2011;40(4, Part 2):e219–e235.

11. Callender LA, Carroll EC, Beal RWJ, et al. Human CD8+ EMRA T cells display a senescence-associated secretory phenotype regulated by p38 MAPK. 10.1111/acel.12675. Aging Cell. 2018/02/01 2018;17(1):e12675.

12. Benveniste O, Allenbach Y. Inclusion body myositis: accumulation of evidence for its autoimmune origin. Brain. 2019;142(9):2549–2551.

13. Rigolet M, Hou C, Baba Amer Y, et al. Distinct interferon signatures stratify inflammatory and dysimmune myopathies. RMD open. 2019;5(1):e000811.

14. Yamashita S, Tawara N, Zhang Z, et al. Pathogenic role of anti-cN1A autoantibodies in sporadic inclusion body myositis. J Neurol Neurosurg Psychiatry. Dec 2023;94(12):1018–1024.

15. Lilleker JB, Naddaf E, Saris CGJ, Schmidt J, de Visser M, Weihl CC. 272nd ENMC international workshop: 10 Years of progress - revision of the ENMC 2013 diagnostic criteria for inclusion body myositis and clinical trial readiness. 16-18 June 2023, Hoofddorp, The Netherlands. Neuromuscul Disord. Apr 2024;37:36–51.

16. Preuße C, Goebel HH, Held J, et al. Immune-mediated necrotizing myopathy is characterized by a specific Th1-M1 polarized immune profile. The American journal of pathology. Dec 2012;181(6):2161–71.

17. Preuße C, Allenbach Y, Hoffmann O, et al. Differential roles of hypoxia and innate immunity in juvenile and adult dermatomyositis. Acta Neuropathologica Communications. 2016/04/27 2016;4(1):45.

18. Udd B, Stenzel W, Oldfors A, et al. 1st ENMC European meeting: The EURO-NMD pathology working group Recommended Standards for Muscle Pathology Amsterdam, The Netherlands, 7 December 2018. Neuromuscular disorders : NMD. Jun 2019;29(6):483–485.

19. Kleefeld F, Horvath R, Pinal-Fernandez I, et al. Multi-level profiling unravels mitochondrial dysfunction in myotonic dystrophy type 2. Acta Neuropathol. Jan 19 2024;147(1):19.

20. Gangfuß A, Hentschel A, Heil L, et al. Proteomic and morphological insights and clinical presentation of two young patients with novel mutations of BVES (POPDC1). Mol Genet Metab. Jul 2022;136(3):226–237.

21. Ladislau L, Suárez-Calvet X, Toquet S, et al. JAK inhibitor improves type I interferon induced damage: proof of concept in dermatomyositis. Brain. Jun 1 2018;141(6):1609–1621.

22. Perez-Riverol Y, Bai J, Bandla C, et al. The PRIDE database resources in 2022: a hub for mass spectrometry-based proteomics evidences. Nucleic Acids Res. Jan 7 2022;50(D1):D543–d552.

23. Hathazi D, Griffin H, Jennings MJ, et al. Metabolic shift underlies recovery in reversible infantile respiratory chain deficiency. The EMBO Journal. 2020;39(23):e105364.

24. Keraite I, Becker P, Canevazzi D, et al. A method for multiplexed full-length single-molecule sequencing of the human mitochondrial genome. Nat Commun. Oct 6 2022;13(1):5902.

25. Pinal-Fernandez I, Casal-Dominguez M, Derfoul A, et al. Machine learning algorithms reveal unique gene expression profiles in muscle biopsies from patients with different types of myositis. Annals of the rheumatic diseases. Sep 2020;79(9):1234–1242.

26. Kleefeld F, Uruha A, Schänzer A, et al. Morphologic and Molecular Patterns of Polymyositis With Mitochondrial Pathology and Inclusion Body Myositis. Neurology. Nov 15 2022;99(20):e2212–e2222.

27. Fayet G, Jansson M, Sternberg D, et al. Ageing muscle: clonal expansions of mitochondrial DNA point mutations and deletions cause focal impairment of mitochondrial function. Neuromuscular disorders : NMD. Jun 2002;12(5):484–93.

28. Carpenter S. Inclusion body myositis, a review. Journal of neuropathology and experimental neurology. Nov 1996;55(11):1105–14.

29. Chen T, He J, Huang Y, Zhao W. The generation of mitochondrial DNA large-scale deletions in human cells. Journal of Human Genetics. 2011/10/01 2011;56(10):689–694.

30. Dong J, Chen L, Ye F, et al. Mic19 depletion impairs endoplasmic reticulum-mitochondrial contacts and mitochondrial lipid metabolism and triggers liver disease. Nature Communications. 2024/01/02 2024;15(1):168.

31. Kim J, Kim HS, Chung JH. Molecular mechanisms of mitochondrial DNA release and activation of the cGAS-STING pathway. Exp Mol Med. Mar 2023;55(3):510–519.

32. Tumburu L, Ghosh-Choudhary S, Seifuddin FT, et al. Circulating mitochondrial DNA is a proinflammatory DAMP in sickle cell disease. Blood. Jun 3 2021;137(22):3116–3126.

33. Alhatou MI, Sladky JT, Bagasra O, Glass JD. Mitochondrial abnormalities in dermatomyositis: characteristic pattern of neuropathology. J Mol Histol. Aug 2004;35(6):615–9.

34. Schänzer A, Rager L, Dahlhaus I, et al. Morphological Characteristics of Idiopathic Inflammatory Myopathies in Juvenile Patients. Cells. 2022;11(1):109.

35. Hedberg-Oldfors C, Lindgren U, Visuttijai K, et al. Respiratory chain dysfunction in perifascicular muscle fibres in patients with dermatomyositis is associated with mitochondrial DNA depletion. Neuropathol Appl Neurobiol. Dec 2022;48(7):e12841.

36. Meyer A, Laverny G, Allenbach Y, et al. IFN-β-induced reactive oxygen species and mitochondrial damage contribute to muscle impairment and inflammation maintenance in dermatomyositis. Acta Neuropathologica. 2017/10/01 2017;134(4):655–666.

37. Oldfors A, Moslemi AR, Jonasson L, Ohlsson M, Kollberg G, Lindberg C. Mitochondrial abnormalities in inclusion-body myositis. Neurology. 2006;66(1 suppl 1):S49–S55.

38. Bhatt PS, Tzoulis C, Balafkan N, et al. Mitochondrial DNA depletion in sporadic inclusion body myositis. Neuromuscular disorders : NMD. Mar 2019;29(3):242–246.

39. Oikawa Y, Izumi R, Koide M, et al. Mitochondrial dysfunction underlying sporadic inclusion body myositis is ameliorated by the mitochondrial homing drug MA-5. PLoS One. 2020;15(12):e0231064.

40. Askanas V, Engel WK, Nogalska A. Sporadic inclusion-body myositis: A degenerative muscle disease associated with aging, impaired muscle protein homeostasis and abnormal mitophagy. Biochimica et Biophysica Acta (BBA) - Molecular Basis of Disease. 2015/04/01/ 2015;1852(4):633–643.

41. Vincent AE, Ng YS, White K, et al. The Spectrum of Mitochondrial Ultrastructural Defects in Mitochondrial Myopathy. Sci Rep. Aug 10 2016;6:30610.

42. Marchi S, Guilbaud E, Tait SWG, Yamazaki T, Galluzzi L. Mitochondrial control of inflammation. Nat Rev Immunol. Mar 2023;23(3):159–173.

43. Picca A, Faitg J, Auwerx J, Ferrucci L, D’Amico D. Mitophagy in human health, ageing and disease. Nat Metab. Dec 2023;5(12):2047–2061.

44. Sachdev R, Kappes-Horn K, Paulsen L, et al. Endoplasmic Reticulum Stress Induces Myostatin High Molecular Weight Aggregates and Impairs Mature Myostatin Secretion. Mol Neurobiol. Nov 2018;55(11):8355–8373.

45. Pinal-Fernandez I, Casal-Dominguez M, Derfoul A, et al. Identification of distinctive interferon gene signatures in different types of myositis. Neurology. Sep 17 2019;93(12):e1193–e1204.

46. Decout A, Katz JD, Venkatraman S, Ablasser A. The cGAS–STING pathway as a therapeutic target in inflammatory diseases. Nature Reviews Immunology. 2021/09/01 2021;21(9):548–569.

47. Britson KA, Ling JP, Braunstein KE, et al. Loss of TDP-43 function and rimmed vacuoles persist after T cell depletion in a xenograft model of sporadic inclusion body myositis. Sci Transl Med. Jan 19 2022;14(628):eabi9196.

48. Rygiel KA, Miller J, Grady JP, Rocha MC, Taylor RW, Turnbull DM. Mitochondrial and inflammatory changes in sporadic inclusion body myositis. 10.1111/nan.12149. Neuropathology and Applied Neurobiology. 2015/04/01 2015;41(3):288–303.

49. Lindgren U, Roos S, Hedberg Oldfors C, Moslemi AR, Lindberg C, Oldfors A. Mitochondrial pathology in inclusion body myositis. Neuromuscul Disord. Apr 2015;25(4):281–8.

50. Yu CH, Davidson S, Harapas CR, et al. TDP-43 Triggers Mitochondrial DNA Release via mPTP to Activate cGAS/STING in ALS. Cell. Oct 29 2020;183(3):636–649.e18.

51. Olivé M, Janué A, Moreno D, Gámez J, Torrejón-Escribano B, Ferrer I. TAR DNA-Binding protein 43 accumulation in protein aggregate myopathies. Journal of neuropathology and experimental neurology. Mar 2009;68(3):262–73.

52. Cavalcante WCP, da Silva AMS, Mendonça RH, et al. Whole-body muscle magnetic resonance imaging in inflammatory myopathy with mitochondrial pathology. Front Neurol. 2024;15:1386293.

53. Zierer LK, Naegel S, Schneider I, et al. Quantitative whole-body muscle MRI in idiopathic inflammatory myopathies including polymyositis with mitochondrial pathology: indications for a disease spectrum. J Neurol. Jun 2024;271(6):3186–3202.

