## Supplementary figures and images for "Mitochondrial leakage and mtDNA damage trigger early immune response in Inclusion Body Myositis"

### Supplementary Fig 1

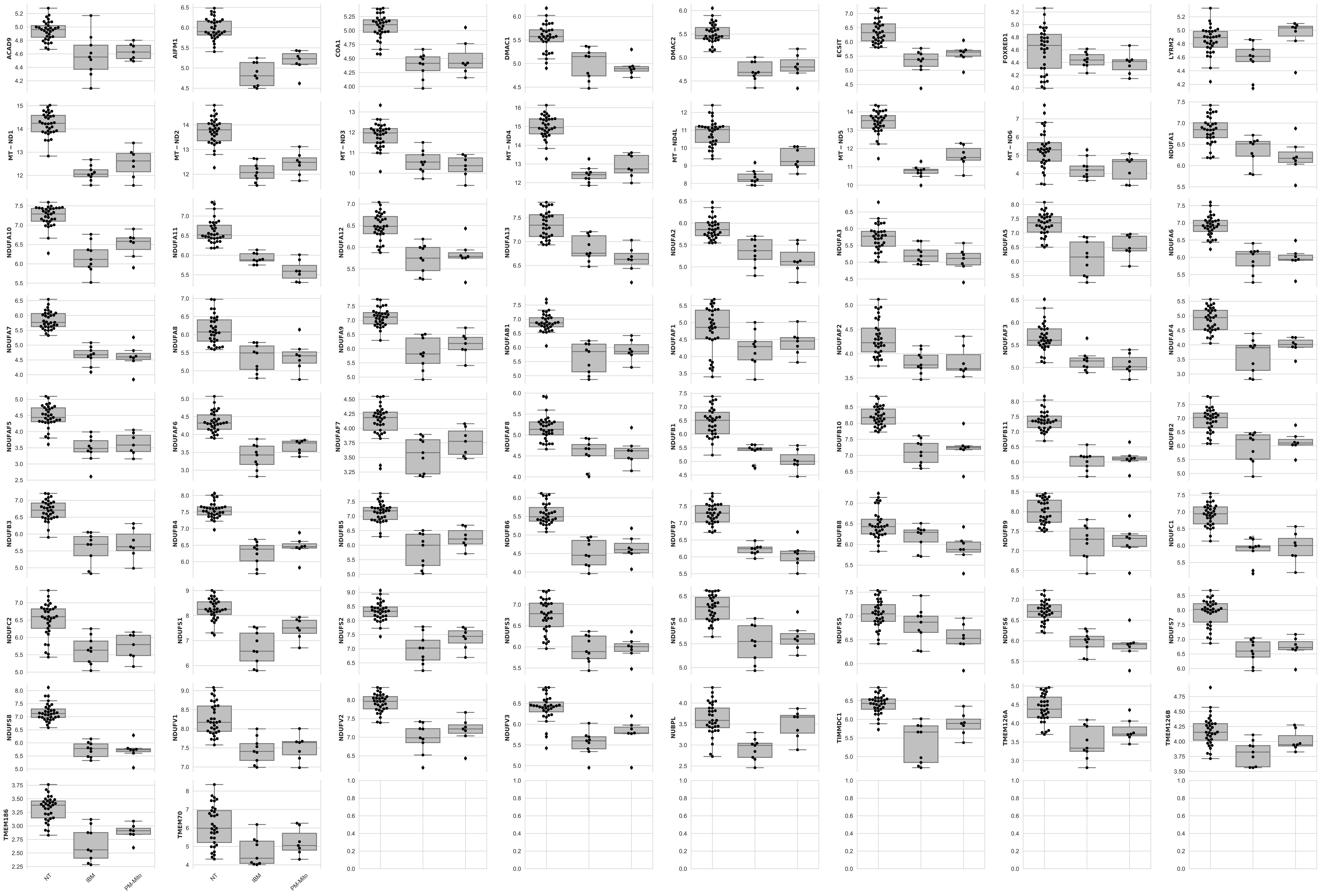

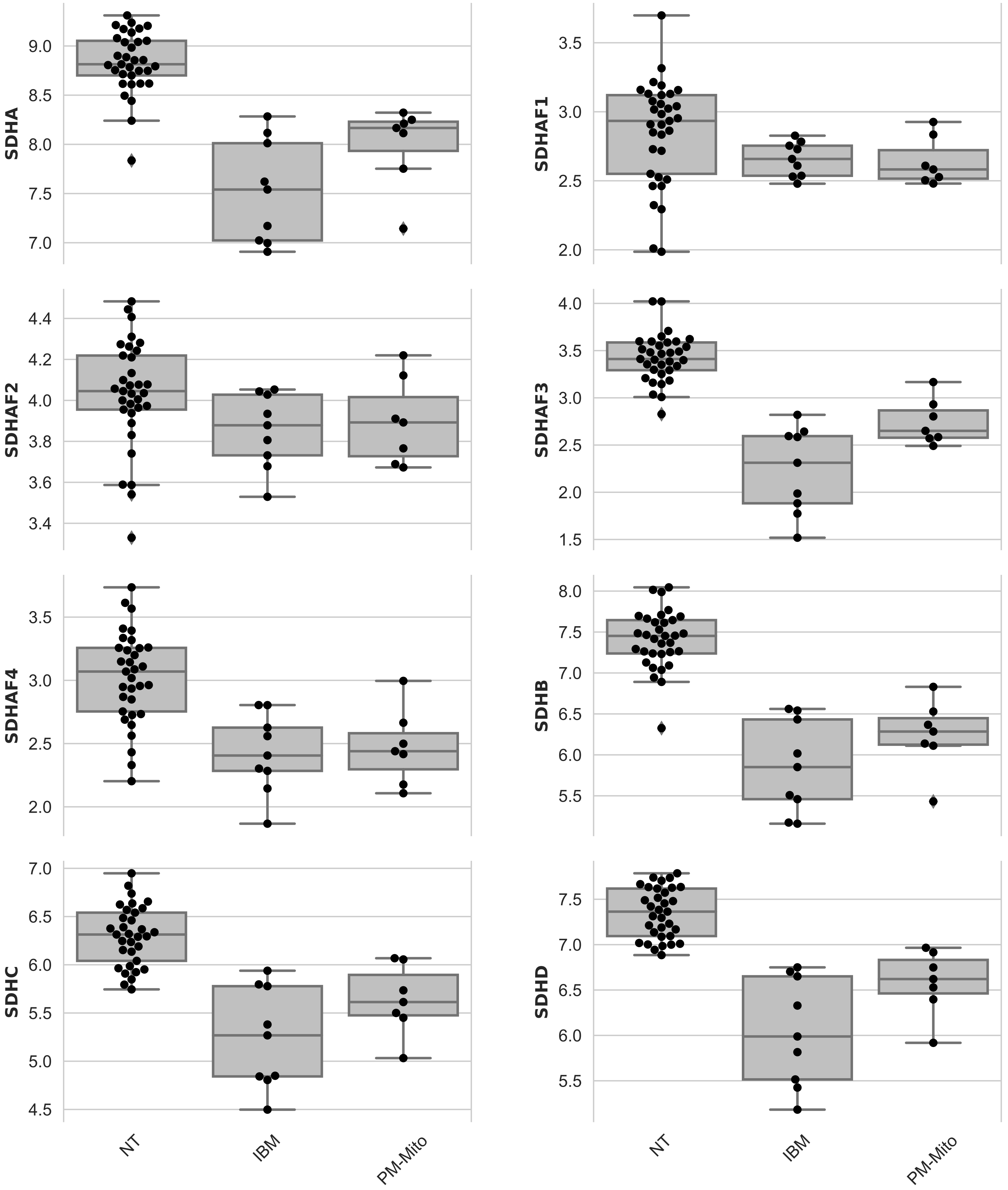

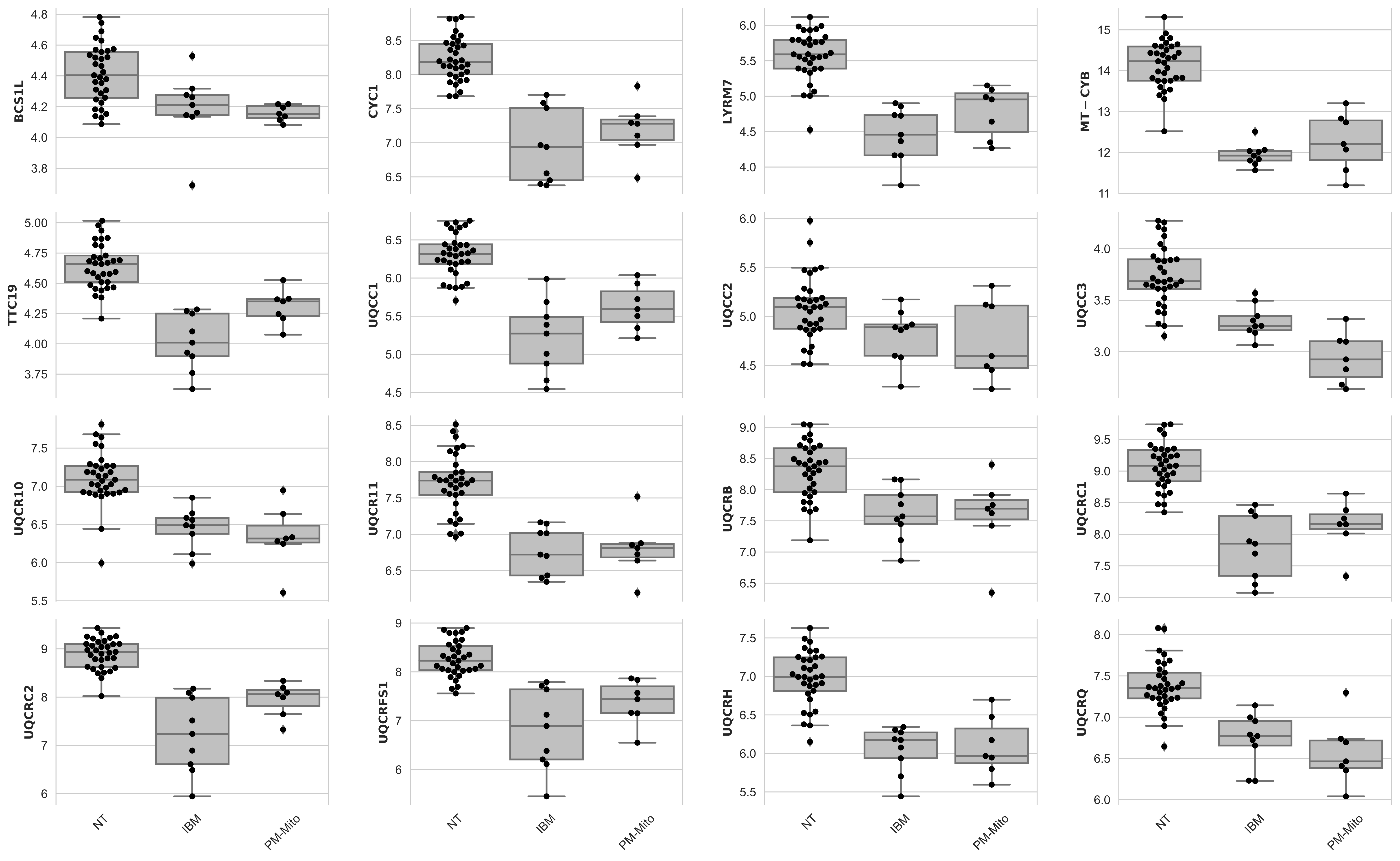

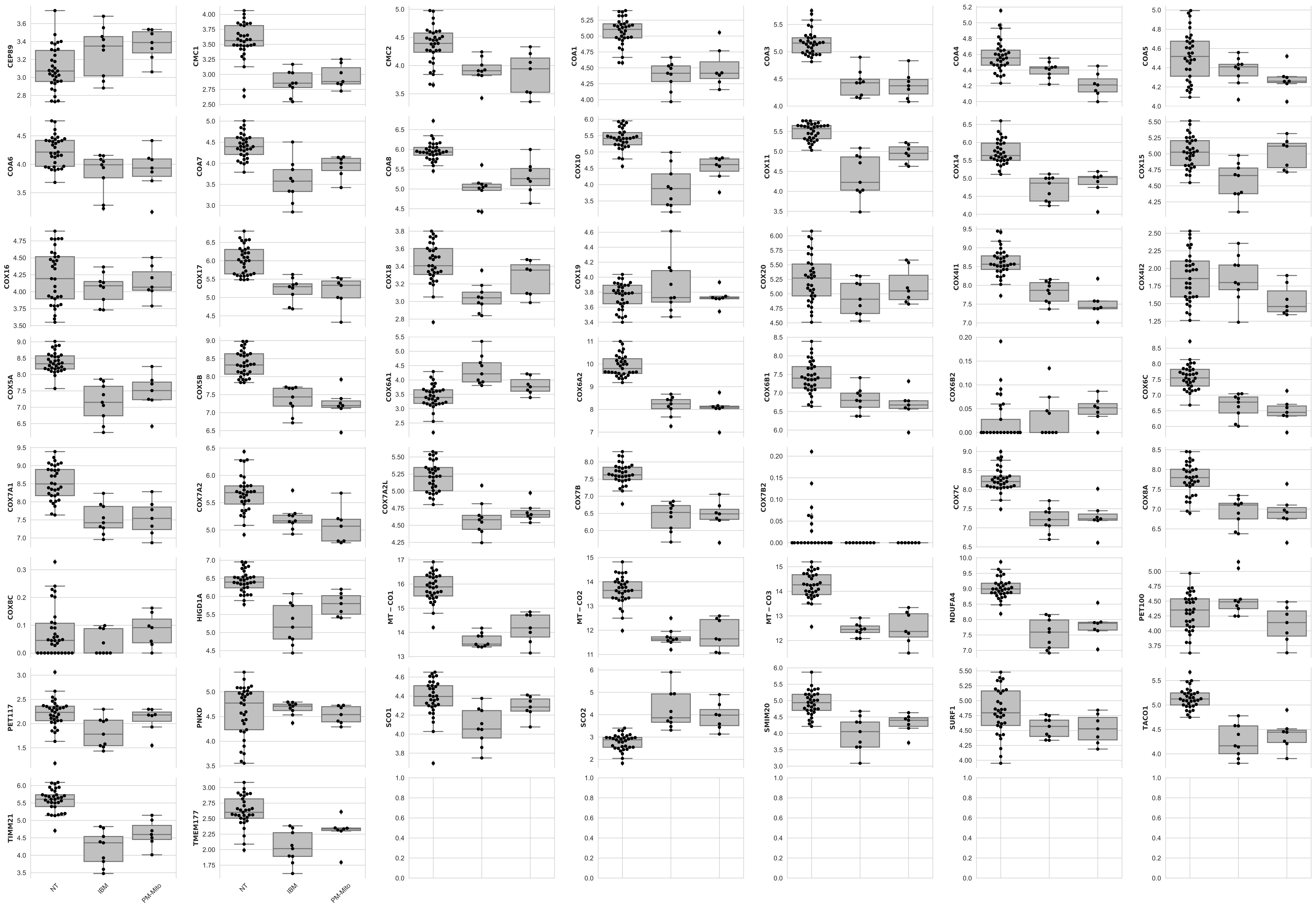

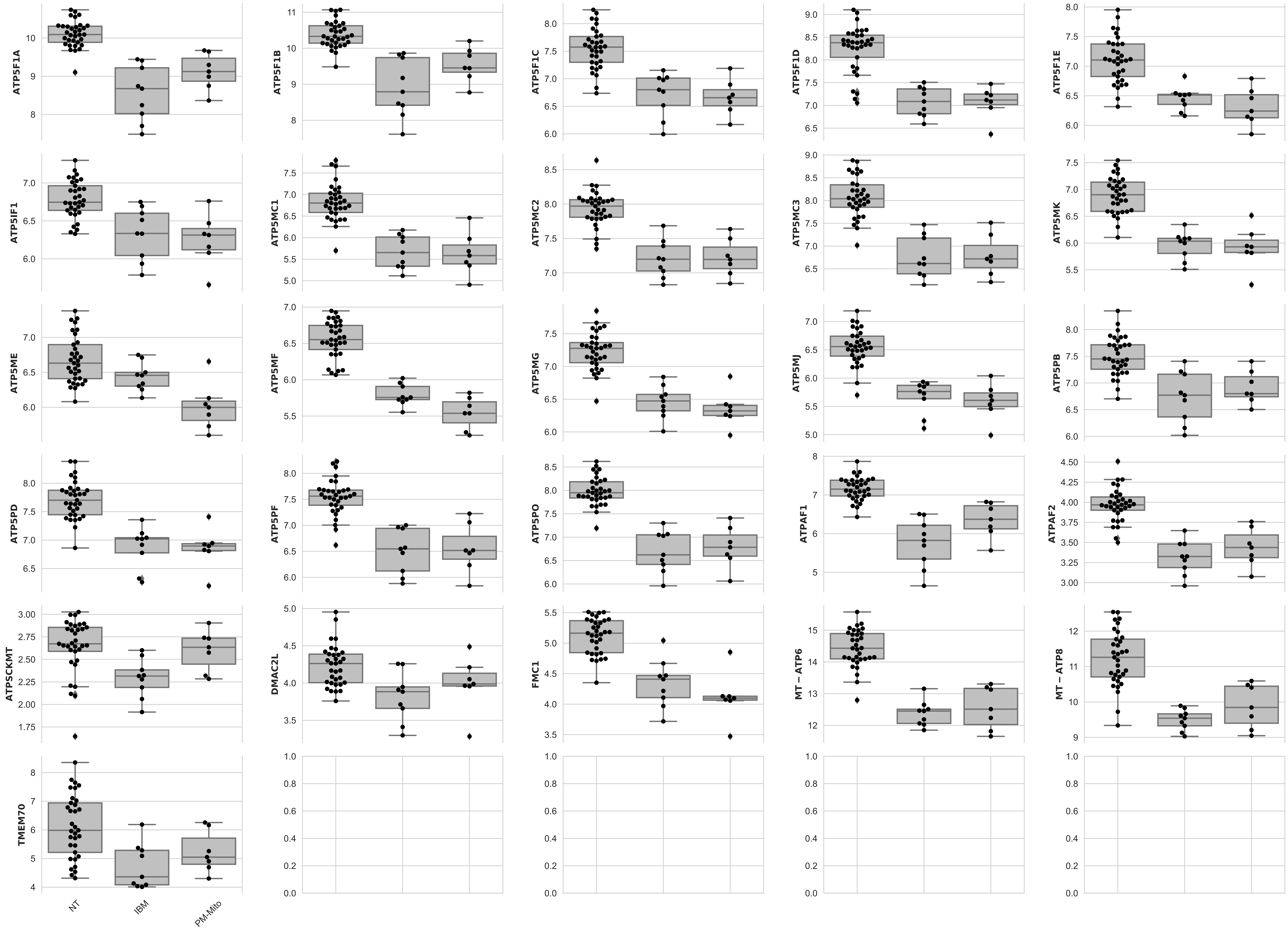

### Supplementary Fig 2

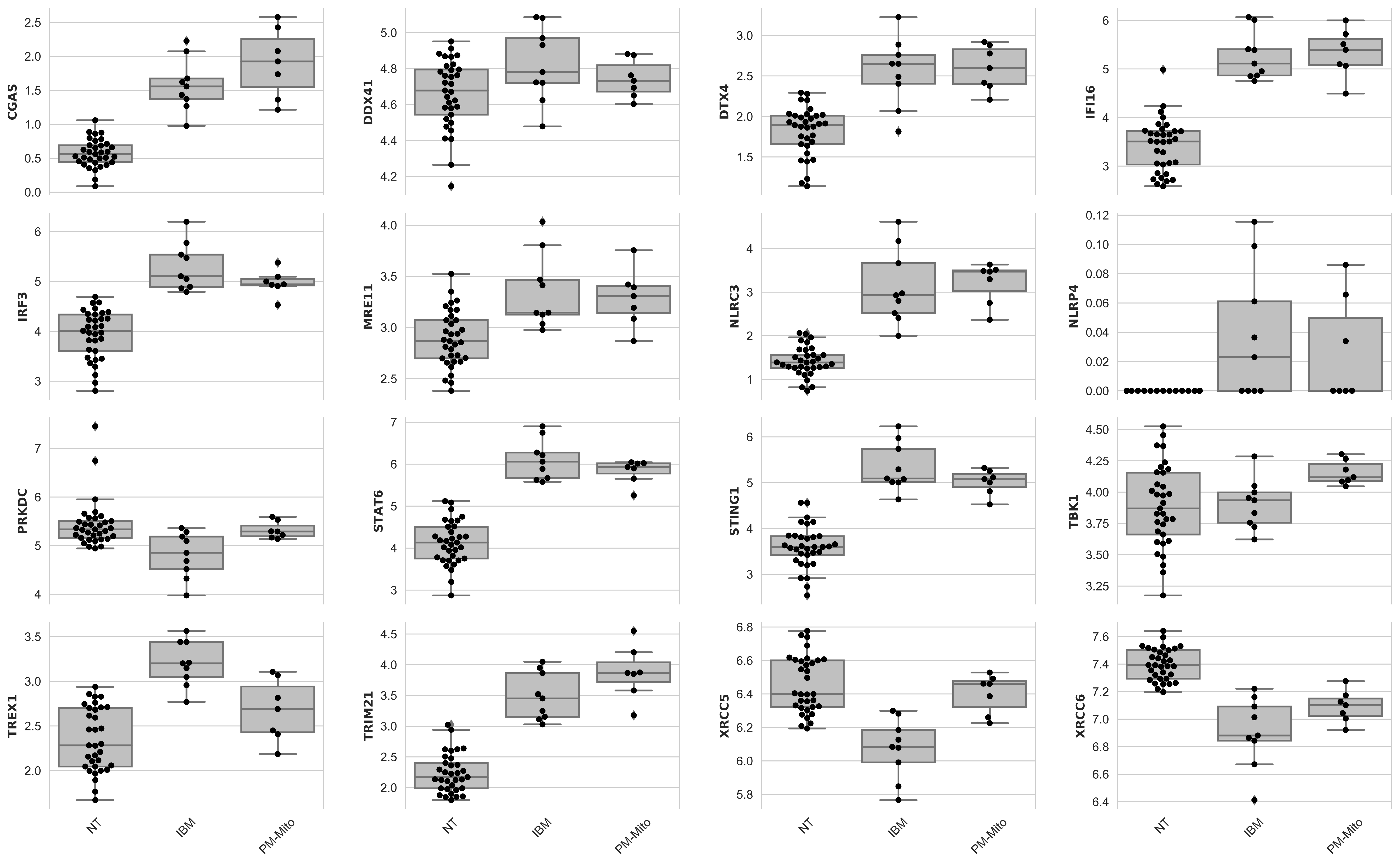

### Supplementary Fig 3

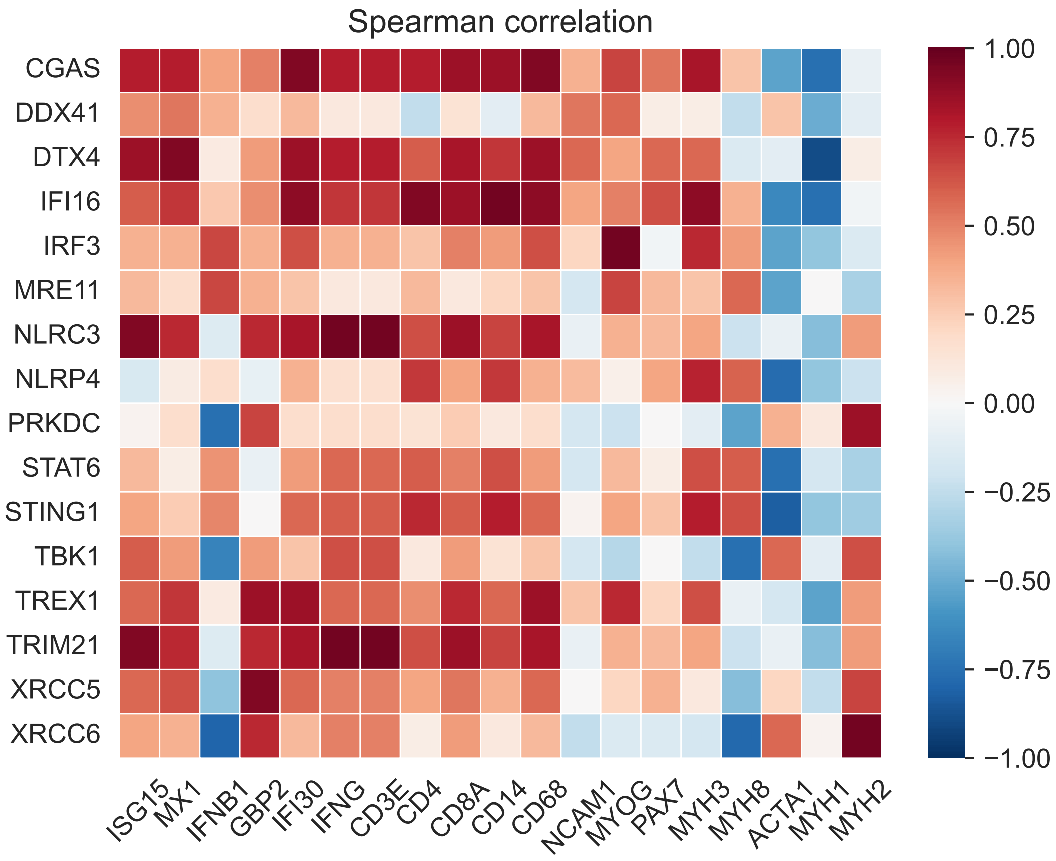
